# Chromosome folding and prophage activation reveal gut-specific genome dynamics of bacteria in the OMM^12^ consortium

**DOI:** 10.1101/2022.05.18.492453

**Authors:** Quentin Lamy-Besnier, Amaury Bignaud, Julian R. Garneau, Marie Titecat, Devon Conti, Alexandra Von Strempel, Marc Monot, Bärbel Stecher, Romain Koszul, Laurent Debarbieux, Martial Marbouty

## Abstract

Bacteria and their viruses, bacteriophages, are the most abundant entities of the gut microbiota, a complex community of microorganisms associated with human health and disease. In this ecosystem the interactions between these two key components are still largely unknown. In particular, the impact of the gut environment on bacteria and their associated prophages is yet to be deciphered. To gain insight into the activity of lysogenic phages within the context of their host genomes, we performed Hi-C on the 12 strains of the OMM^12^ synthetic bacterial community stably associated within mice gut (gnotobiotic mouse line OMM^12^) in both *in vitro* and *in vivo* conditions. High-resolution contact maps of the chromosome 3D organization of the bacterial genomes revealed a wide diversity of architectures, differences between environments and an overall stability over time in the gut of mice. The DNA contacts also pointed at 3D signatures of prophages leading to predict 16 of them as functional. We identified circularization signals and observed different 3D patterns depending on the condition. Concurrent virome analysis showed that 11 of these prophages produced viral particles *in vivo* and/or *in vitro*, and that OMM^12^ mice do not carry other intestinal viruses. By predicting functional prophages, the Hi-C approach unlocks the study of phage-bacteria interaction dynamics.

## INTRODUCTION

Microbial communities that inhabit the mammal intestinal tract are complex and predominantly composed of a variety of bacteria and their specific viruses, the bacteriophages (phages). Variations in the composition of both bacterial and viral intestinal communities have been associated with a wide range of diseases (Mangalea et al., 2021; Norman et al., 2015; Yang et al., 2021; Zhao et al., 2017). However, the mechanisms supporting these associations are still poorly understood. Indeed, the broad diversity of intestinal microbes residing in the gut of humans or laboratory animals hinder the precise identification of the role of each component.

Over the last decade, multiple studies have employed proximity ligation-based technologies (Dekker et al., 2002) (e.g. Hi-C) to study the 3D organization of bacterial genomes (Le et al., 2013a; Lioy et al., 2018; Marbouty et al., 2015; Umbarger et al., 2011; Val et al., 2016; Wang et al., 2015). These approaches, which quantify the relative collision frequencies between DNA segments, revealed local and global folding of bacterial chromosomes. Using various growth conditions and mutants in combination with other omics approaches, these studies have allowed to better understand the role of nucleoid-associated proteins (NAPs) and transcription in the 3D genome architecture of model bacteria (Le and Laub, 2016; Lioy et al., 2018; Marbouty et al., 2015). However, major clades of the intestinal microbial communities remain unexplored. Moreover, most studies have focused on *in vitro* cultures leaving open questions on the 3D chromosomal architectures in the digestive tract.

In parallel, we and others have developed Hi-C derivatives approaches to improve metagenome analysis (e.g. meta3C or metaHi-C) (Beitel et al., 2014; Burton, 2014; Marbouty et al., 2014) (reviewed in (Marbouty and Koszul, 2015)). Taking advantage of DNA contacts as a marker of relative physical proximity, we and others developed and applied metaHi-C to successfully infer the bacterial hosts of episomes, such as plasmids and phages, in complex natural microbial populations (Bickhart et al., 2019; Marbouty et al., 2014, 2017, 2021; Stalder et al., 2019). We also demonstrated the relevance of using Hi-C data to study DNA segment integration in bacterial genomes as well as the physiological state of prophages through detection of typical 3D signatures (Marbouty et al., 2014, 2015, 2017).

Gnotobiotic animals, in which a defined population of microbes is introduced, provide a path to a reductionist approach. In 2016, the mouse line named Oligo-MM12 (OMM^12^), was proposed to the community as a platform from which the role of individual bacteria and environmental parameters (e.g. diet, infection) could be investigated with high reproducibility (Brugiroux et al., 2016). These 12 strains were initially isolated from conventional mice intestines and are representative of the five most abundant phyla in the mouse gut microbiota: *Bacteroidetes, Firmicutes, Actinobacteria, Proteobacteria* and *Verrucomicrobia*. Comparison of four different breeding facilities demonstrated negligible variations in the composition and relative abundance of these gut species (Eberl et al., 2020). Another advantage of this model is the possibility to track the genomic information over time as the 12 strains have been sequenced and assembled (Garzetti et al., 2017; Lamy-Besnier et al., 2021). Furthermore, additional bacteria can be engrafted without disrupting the resident microbiota to eventually provide additive functions (Bolsega et al., 2019; Brugiroux et al., 2016; Herp et al., 2019; Lourenço et al., 2020; Streidl et al., 2021).

In the present study, we combined Hi-C with virome sequencing to characterize the 3D organization and behavior of chromosomes of bacteria of the OMM^12^ consortium as well as their associated (pro)phages *in vitro* but also *in vivo* (*i*.*e*. in the gut environment). Hi-C contact maps of individual bacteria (*in vitro*) reinforce the central role played by the ParABS system in bacterial chromosome folding and revealed a diversity of architectures found in those bacteria. Comparison with genome-wide contact maps of the same bacteria obtained from *in vivo* gut conditions revealed that the overall chromosome folding is conserved with, nonetheless, some notable differences. As hinted previously (Marbouty et al., 2017), a careful analysis of the data allowed us to predict 16 functional prophages out of 44 potential prophages predicted bioinformatically, to refine the coordinates of their borders, identify circularization events as well as their activity status. 11 of these 16 prophages were confirmed to be active by virome approach that detected a total of 13 induced prophages *in vitro* and / or *in vivo*. These analyses also emphasize that some differences arose between the induction of the prophages in the two studied conditions, highlighting the influence of the gut environment on phage induction and microbial community in general. The same analysis performed one year apart in the same breeding facility revealed that the prophages were induced in a similar way, showing that they constitute a resident phages population in OMM^12^ mice gut. Finally, we were not able to identify any ssDNA or strictly lytic phage in this community.

Our results demonstrate that Hi-C is a compelling resource for studying controlled communities such as OMM^12^ gut, informing on the genome architecture of individual bacteria and predicting functional prophages as well as their activities.

## RESULTS

### *In vitro* characterization of chromosome folding in individual OMM^12^ bacteria revealed common principles of bacteria genomes architectures

We applied a recently published high-resolution version of the Hi-C protocol adapted to bacteria to generate genome-wide contact maps for each of the 12 bacterial strains of the OMM^12^ consortium (Fig. 1, Supplementary Fig. 1 and Supplementary Table 1) (Methods, (Cockram et al., 2020)). In model bacteria species, these contact maps typically display shared characteristics. First, a strong and broad diagonal reflecting frequent local contacts between neighboring loci. This property can be exploited to scaffold incomplete genomes (reviewed in Flot et al., 2015), and was recently applied to close nine genomes of the OMM^12^ consortium (Lamy-Besnier et al., 2021). Three genomes of the consortium (*B. caecimuris, B. animalis* and *F. plautii*) remained nevertheless incomplete, and were therefore scaffolded using the last published sequences (Supplementary Fig. 2). Next, bacterial contact maps typically exhibit self-interacting domains (“chromatin interaction domains” or CIDs) visualized as squares along the main diagonal (Le et al., 2013a; Lioy et al., 2018; Marbouty et al., 2015; Umbarger et al., 2011; Val et al., 2016), whose boundaries correlate with high transcriptional activity (Le et al., 2013a; Lioy et al., 2018). As expected, CIDs were also found in the 12 contact maps, ranging in size from tens to hundreds of kilobases as previously observed for other bacteria (Supplementary Fig. 1) and with many boundaries colocalizing with transcriptionally active rRNA or tRNA loci. In addition, most bacteria studied so far (with the notable exception of *E. coli* (Lioy et al., 2018) and to a lesser extent *V. cholerae* (Val et al., 2016)) display an anti-diagonal perpendicular to the main one and extending from the origin of replication down to the terminus (Böhm et al., 2020; Marbouty et al., 2015; Umbarger et al., 2011). This pattern, which reflects enriched contacts between the two replichores along their entire length, was clearly observed for 7 bacteria of the OMM^12^ (*B. caecimuris, T. muris, A. muris, E. clostridioformis, F. plautii, B. coccoides, C. innocuum*), barely visible in 2 (*A. muciniphila, E. faecalis*) and not detectable in the 3 others (*M. intestinale, B. longum, L. reuteri*) (Fig. 1 and Supplementary Fig. 1). Interestingly, the different matrices generated also revealed the diversity of bacteria chromosomes architectures with various structures such as the bow shape signals observed in *B. coccoides* and *L. reuteri* contact map. The 12 matrices generated in this experiment confirm common principles but also reveal a large diversity of bacteria chromosome folding.

**Table 1:** Candidate prophages of the OMM^12^ consortium. The localization and size of the putative prophages as predicted by Vibrant and Virsorter are indicated. When Vibrant and Virsorter disagreed, the combined longest location was selected. In the localization column bold face indicates induced prophages described by Zund *et al*.. Detected 3D patterns as well as their variations between *in vitro* and *in vivo* conditions are indicated (dark grey = strong signal, light grey = week signal). NA: Not applicable (coverage too low). Induced phages detected through virome sequencing are indicated in the last column.

| Bacteria |  | Localisation (Mbp) | Vibrant | Virsorter2 | 3C in vitro | 3C in vivo | 3C in vitro vs in vivo | Virome |
| --- | --- | --- | --- | --- | --- | --- | --- | --- |
| *A. muris* | **KB18** | 0.026-0.095 |  | Yes (Cat6) |  |  |  |  |
|  |  | **1.053-1.120** |  | Yes (Cat5) | Yes (TAD) | NA | NA | Yes |
|  |  | 2.999-3.151 | Yes | Yes (Cat6) | Yes (TAD) | NA | NA |  |
|  |  | 3.228-3.310 |  | Yes (Cat5) |  |  |  |  |
| *A. muciniphila* | **YL44** | 0.038-0.072 |  | Yes (Cat6) | Yes (TAD) | Yes (TAD) | same |  |
|  |  | **0.712-0.743** | Yes | Yes (Cat5) | Yes (TAD) | Yes (TAD) | same | Yes |
|  |  | 1.336-1.370 | Yes | Yes (Cat6) |  |  |  | Yes |
|  |  | **2.291-2.337** | Yes | Yes (Cat5) | Yes (depletion) | Yes (depletion) | same | Yes |
| *B. caecimuris* | **I48** | 0.271-0.355 |  | Yes (Cat6) |  |  |  |  |
|  |  | 0.922-1.008 |  | Yes (Cat6) |  | Yes (TAD + loop) |  |  |
|  |  | 1.409-1.421 |  | Yes (Cat6) |  |  |  |  |
|  |  | 2.969-3.118 | Yes | Yes (Cat4) | Yes (loop + depletion) | Yes (TAD + loop) | increased coverage and signal | Yes |
| *B. coccoides* | **YL58** | 1.287-1.351 |  | Yes (Cat5) |  |  |  |  |
|  |  | 1.845-1.851 | Yes |  |  |  |  |  |
|  |  | 2.228-2.268 |  | Yes (Cat6) |  |  |  |  |
|  |  | **3.571-3.620** | Yes | Yes (Cat6) | Yes (TAD + loop) | Yes (TAD + loop) | same | Yes |
|  |  | 4.091-4.112 | Yes |  |  |  |  |  |
| *E. clostridioformis* | **YL32** | 1.254-1.385 | Yes |  |  |  |  |  |
|  |  | 1.275-1.461 |  | Yes (Cat5) |  |  |  |  |
|  |  | 1.398-1.461 | Yes |  |  |  |  |  |
|  |  | 1.613-1.669 | Yes | Yes (Cat5) |  |  |  |  |
|  |  | 2.059-2.106 | Yes | Yes (Cat4) | Yes (TAD + depletion signal) | Yes (TAD + loop) | increased loop signal | Yes |
|  |  | 2.326-2.428 |  | Yes (Cat6) |  |  |  |  |
|  |  | 2.914-3.014 | Yes | Yes (Cat5) |  |  |  |  |
|  |  | 3.355-3.418 | Yes | Yes (Cat5) | Yes (TAD) | Yes (TAD) | decreased signal | Yes |
|  |  | 4.054-4.122 |  | Yes (Cat6) |  |  |  |  |
|  |  | 4.983-5.030 | Yes |  |  |  |  |  |
|  |  | 5.653-5.746 |  | Yes (Cat6) |  |  |  |  |
|  |  | 6.292-6.376 |  | Yes (Cat6) |  |  |  |  |
|  |  | 6.752-6.852 |  | Yes (Cat6) |  |  |  |  |
| *C. innocuum* | **I46** | 0.454-0.499 |  | Yes (Cat6) | Yes (TAD + loop) | NA | NA |  |
|  |  | 3.744-3.787 | Yes | Yes (Cat5) | Yes (TAD) | NA | NA |  |
|  |  | 4.275-4.452 | Yes | Yes (Cat5) | Yes (multiple TADs) | Yes (TAD) | 2 different phages | Yes |
| *E. faecalis* | **KB18** | 1.672-1.774 |  | Yes (Cat6) |  |  |  |  |
|  |  | 1.972-2.046 |  | Yes (Cat6) |  |  |  |  |
|  |  | 2.899-2.914 | Yes |  |  |  |  |  |
| *F. plautii* | **YL31** | 0.460-0.466 | Yes |  |  |  |  |  |
|  |  | **0.909-1.064** | Yes | Yes (Cat5) | Yes (TAD + loop) | NA | NA | Yes |
|  |  | 1.126-1.212 | Yes | Yes (Cat5) |  |  |  | Yes |
|  |  | **2.738-2.785** | Yes | Yes (Cat5) | Yes (TAD) | NA | NA | Yes |
|  |  | 3.093-3.179 |  | Yes (Cat6) |  |  |  |  |
| *L. reuteri* | **I49** | 0.019-0.045 | Yes |  |  |  |  |  |
| *T. muris* | **YL45** | 2.099-2.135 |  | Yes (Cat6) |  |  |  |  |
|  |  | 2.100-2.144 | Yes | Yes (Cat5) |  |  |  |  |

**Figure 1:**
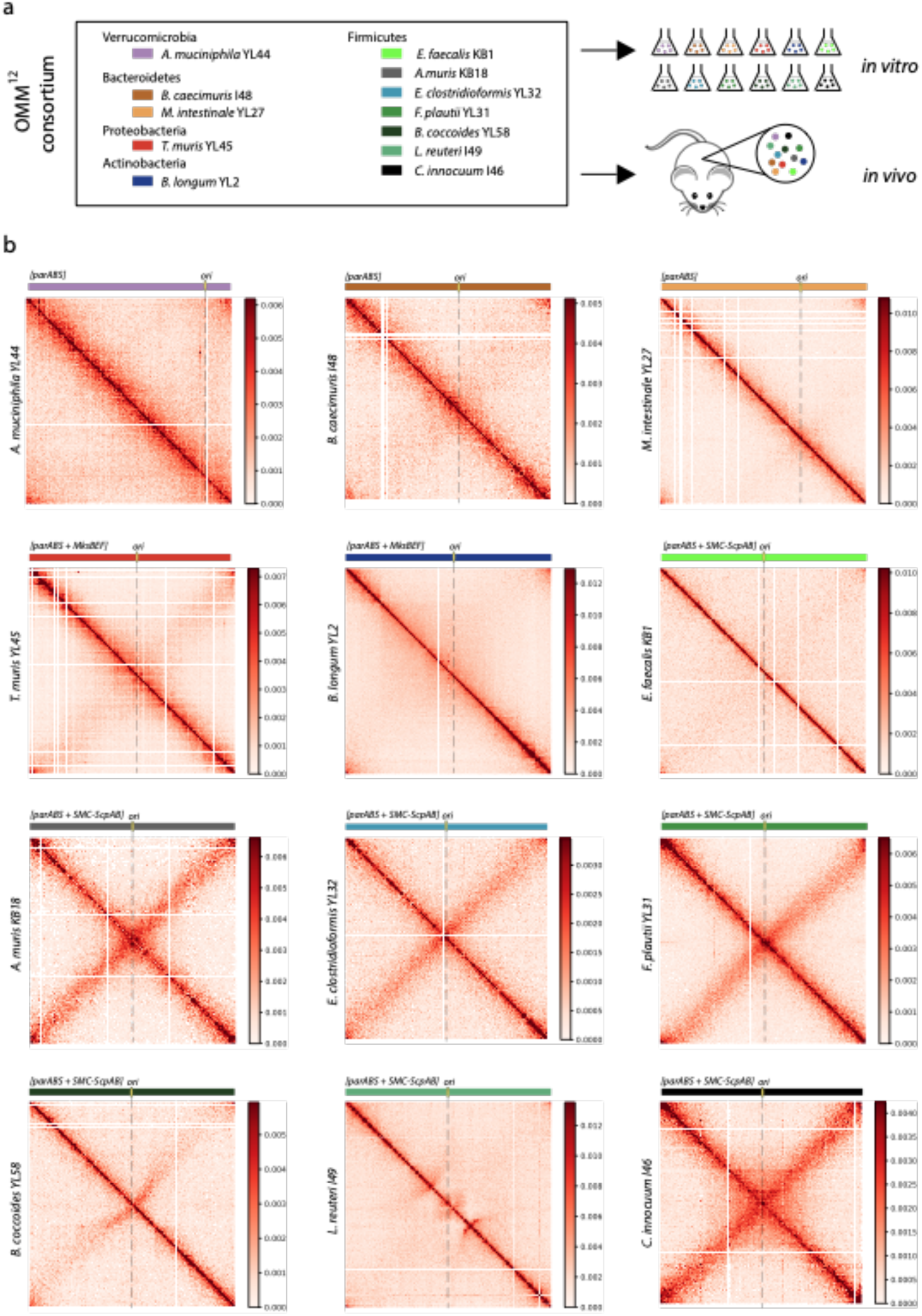
*In vitro* contact maps of the OMM^12^ consortium. **a**. Experimental scheme of the source of Hi-C and virome libraries from the OMM^12^ bacteria. Hi-C and virome were performed both on the independent *in vitro* cultures of each bacterium and from fecal samples. **b**. Contact map obtained from Hi-C performed on *in vitro* cultures for each bacterium of the OMM^12^ consortium (5 kb resolution). The localization of the origin of replication (*ori*) is indicated (black dashed line). Scale bars are indicated aside each matrix.

**Figure 2:**
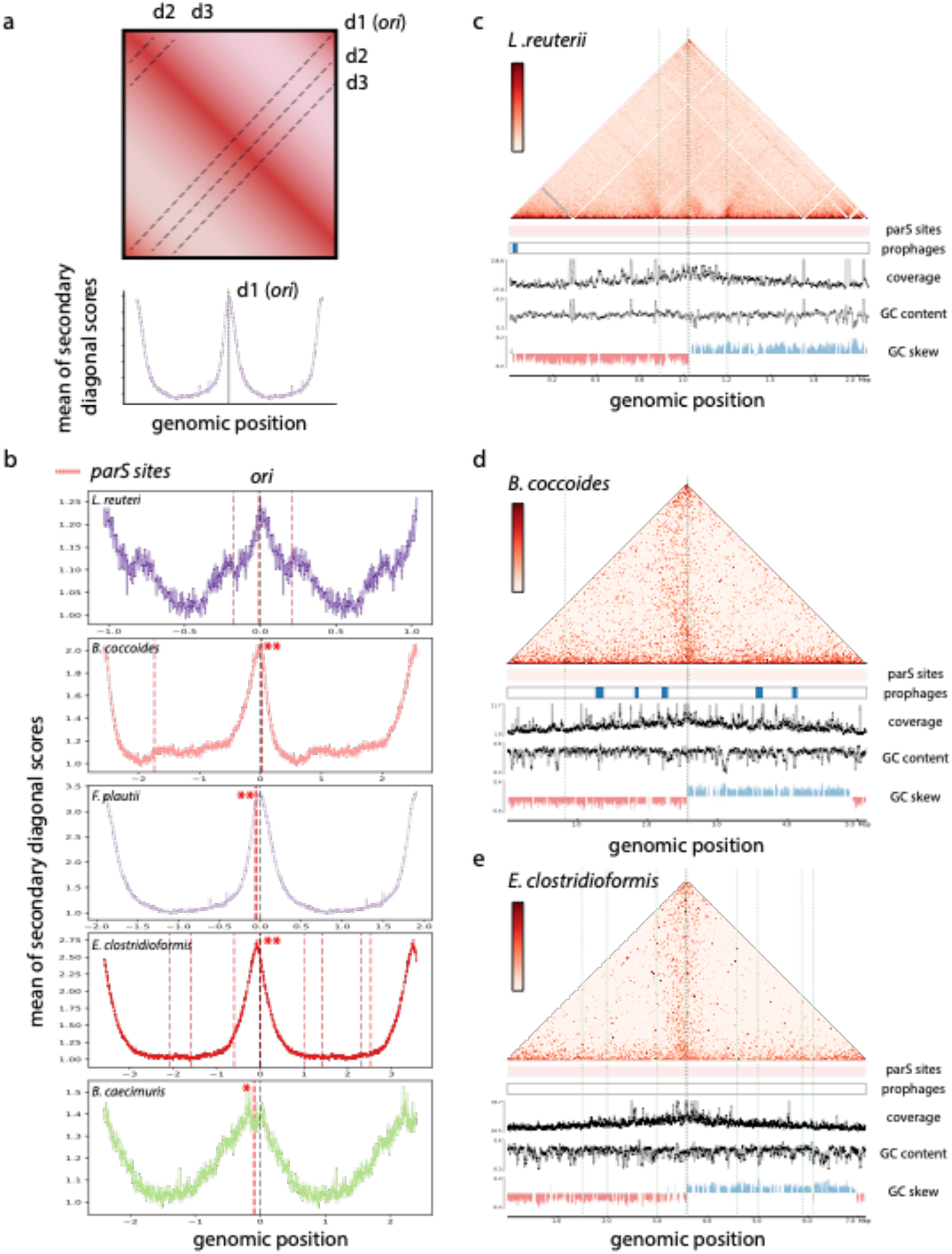
*parS* sites and their implications in the origin domain folding. **a**. Schematic representation of a Hi-C contact map for a bacterial genome. Signal for each secondary diagonal (d1,d2,d3…) was computed in order to generate the graph presented below that showed the strength of the different secondary diagonals along the genome. Each graph was centered on the origin of replication. **b**. Signal of the secondary diagonals for several OMM^12^ bacteria, centered on *ori* (*L. reuteri, B. coccoides, F. plautii, E. clostridioformis, B. caecimuris*). Localisation of *parS* sites are indicated as red dashed lines. Red stars indicate the presence of several *parS* sites in the same 5 kb window. **c-d-e**. Contact map of *L. reuteri* (5 kb resolution), *B. coccoides* (5 kb resolution) and *E. clostridioformis* (5 kb resolution). *ParS* sites (green dashed lines), putative prophages, coverage, GC content, GC skew and genomic coordinates are indicated below each matrix.

### ParS sites as major drivers of ori domains folding

*ParS* sites are widely conserved (Jalal and Le, 2020) in bacteria and previous studies have demonstrated the central role of the ParABS system in regulating the overall 3D organization of bacterial genomes and in the segregation of their new replicated chromosomes (Böhm et al., 2020; Lioy et al., 2020; Marbouty et al., 2015; Wang et al., 2015). The number of *parS* sites varies considerably between bacteria ranging from one to 20 (Livny et al., 2007). Former works have shown that the recruitment of the bacterial structural maintenance of chromosome (SMC) condensin complex SMC-ScpAB at *parS* sites generates hairpins and bridge replichores, through a loop-extrusion like mechanisms (Brandão et al., 2021; Marbouty et al., 2015; Wang et al., 2017). We found homologs of the ParABS system in the 12 genomes confirming its large conservation in bacteria (Methods). Using *parS* consensus sequence, we detected between one and 10 *parS* sites in the 12 genomes with different distributions along the chromosome. In all cases, most of the detected *parS* sites are clustered and positioned at the crossing of the anti-diagonal (Fig. 2 and Supplementary Fig. 3). Using these *parS* clusters and the GC skew, we determined the position of the replication origin on the chromosomes (Methods). The read’s coverage variation supported in all cases these positions. Interestingly, the replication origin does not systematically correlate with the presence of the gene *dnaA*, which is typically used to define it. This suggests caution when using *dnaA* gene in metagenomic studies to characterize contigs encompassing origin of replication (Emiola and Oh, 2018). *ParS* clusters also associate with the presence of a large origin domain (*B. caecimuris, T. muris, A. muris, E. clostridioformis, F. plautii, C. innocuum*) or hairpin structures (*B. coccoides, L. reuteri*) (Supplementary Fig. 1) reminiscent of those observed for *B. subtilis* (Marbouty et al., 2015). We further explore the possible link between *parS* sites, their numbers, their positions and the presence of an anti-diagonal but could not detect any correlation with the strength of the signal in the opposite diagonal (Fig. 2b). The case of *L. reuteri* is particular in that we detect 3 sites that are quite distant to each other but still surround the origin of replication (*parS*1 = -180 kb, *parS*2 = -17 kb, *parS*3 = +206 kb) and, each time, associate with a discrete 3D contact pattern (Fig. 2c). The proximal *parS*2 site is at the center of a small topological domain while the two other sites are associated with the typical hairpin signature observed in *B. subtilis* (Marbouty et al., 2015). This specific structure of origin organization could be the result of the large distance separating the 3 *parS* sites in *L. reuteri*. Our data highlight the impact of *parS* sites distribution in the overall folding of bacterial chromosomes.

**Figure 3:**
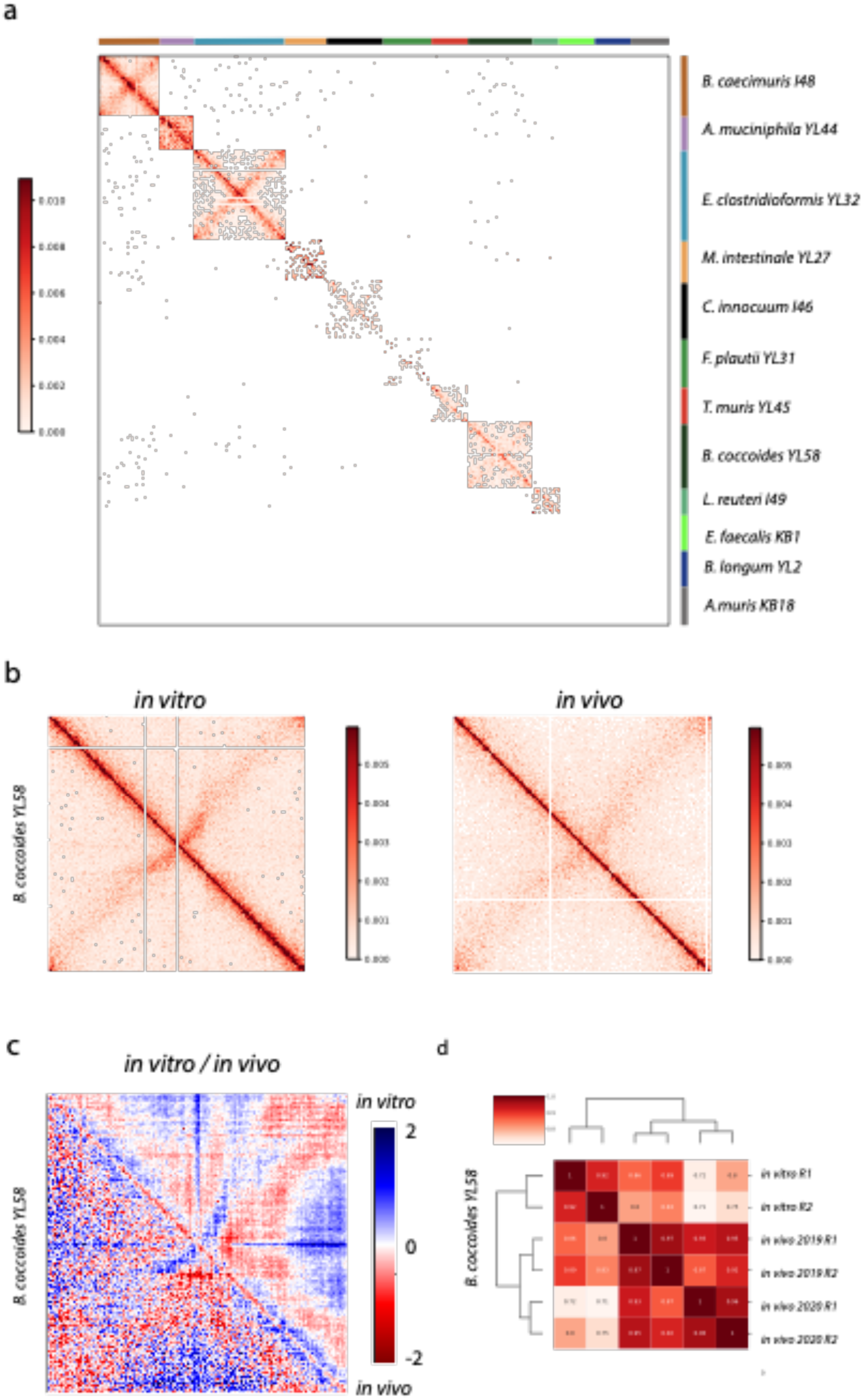
Stability of the chromosome architectures in the gut environment. **a**. Contact map of the entire OMM^12^ consortium (5 kb resolution) obtained from fecal samples. Scale bar is indicated on the left and the different organisms on the right. **b**. Comparison of the contact maps for *B. coccoides* between *in vitro* (left) and *in vivo* (right) conditions. **c**. Ratio (Log2) of contact maps of the two maps of *B. coccoides* shown in panel B. An increase or decrease in contacts is represented in blue (*in vitro)* or red (*in vivo)*, respectively. **d**. Hierarchical clustering of the different Hi-C replicates for *B. coccoides* using the software HiCrep.

### Non-model bacteria rely on yet unidentified SMC complexes for chromosomal arms alignment

Hi-C studies have shed light on the role of the SMC condensin family proteins in maintaining chromosome architecture (Böhm et al., 2020; Le et al., 2013b; Lioy et al., 2020; Marbouty et al., 2015; Wang et al., 2015). Three types of complexes structurally related to condensins have been identified in bacteria: Smc-ScpAB, MukBEF and MksBEF (review in (Nolivos and Sherratt, 2014). Smc-ScpAB is present in most bacteria and often works together with the ParABS system in order to fulfill proper chromosome segregation (Jalal and Le, 2020). In contrast MukBEF is restricted to *Enterobacteria* whereas MksBEF is scattered over the phylogenic tree and both systems do not appear to promote chromosomal arms juxtaposition (Böhm et al., 2020; Lioy et al., 2018, 2020). Among the 12 genomes, seven belonging to the *Firmicutes* phylum contain a clear homolog of the Smc-ScpAB proteins and two (*B. longum* and *T. muris*) have a distant homolog of the MksBEF system. For three bacteria (*A. muciniphila, B. caecimuris* and *M. intestinale*), including the two belonging to the phylum *Bacteroidetes*, no homologs of known bacterial condensins could be found (Methods). This observation contrasts with the idea that this family of proteins is nearly ubiquitous in bacteria (Hirano, 2016; Mäkelä and Sherratt, 2020), and suggest that other distant homologs have yet to be identified. Two of the species encoding Smc-ScpAB homologs do not display an anti-diagonal in their genomic contact maps (*L. reuteri* and *E. faecalis*), indicating that the presence of the Smc-ScpAB system does not systematically lead to a tight bridging of chromosomes arms (Fig. 1 and Fig. 2). On the other hand, we detect in *B. caecimuris* and especially in *T. muris* an anti-diagonal signal despite no homologs of condensins is found in these genomes (Methods). We detect also weak homologs of MksBEF in *T. muris*, a complex that does not promote arm alignment (Böhm et al., 2020).

Our data indicate that possible distant homologs of SMC proteins or even new processes involved in chromosomal arm alignment remain to be uncovered. This Hi-C experiment on non-model bacteria species open new avenues regarding chromosome folding regulation in prokaryotes.

### Bacteria chromosomal architectures present variations but remain stable in the mice gut environment

We next ask whether the chromosome organization of each bacterium grown individually is affected when they are altogether in the gut of mice. We applied our latest metagenomic Hi-C protocol to mouse feces (n=2; September 2019; Methods; (Marbouty et al., 2021)) and generated the resulting contact matrices for the entire consortium and for each bacterium (Fig. 3 and Supplementary Fig. 4). The resulting contact map show no background signal between the different genomes, demonstrating the efficiency of the protocol (Fig. 3a). It also confirms that the genomes are well assembled, without contamination from one bacterium to another. Of the 12 bacteria present in the consortium, only 6 are sufficiently abundant to obtain individual contact matrices with exploitable signal (*B. caecimuris, E. clostridioformis, A. muciniphila, M. intestinale, C. innocum, B. coccoides*). Analysis of the different matrices obtained show that large structures like the anti-diagonal signal were preserved in the intestinal environment while local structures exhibit important differences (Fig. 3b-c and Supplementary Fig. 4). For 3 species (*A. muciniphila, B. caecimuris* and *E. clostridioformis*), we detected an increase of short-range contact in the *in vivo* conditions associated with a decrease of long-range interactions. For *C. innocuum* and *B. coccoides*, we observed an increase of the interactions at very short-range distance in the *in vivo* conditions. Finally, in *M. intestinale*, the contact map from the fecal sample shows notable differences compared to lab conditions with a clear increase of long-range interactions. We also observed a portion of the genome exhibiting multiple long-range interactions, reminiscent of the loops observed in *B. subtilis* (Marbouty et al., 2015) and recently shown to correspond to Rok-dependant contacts (Dugar et al., 2022). We could not detect specific annotations associated with this pattern, and further experiments and analysis will be needed to understand those differences. Our results were confirmed by performing Hi-C on feces sampled 9 months later (n=2; May 2020) from mice bred in the same facility. Comparison using HiCRep ((Yang et al., 2017); Method) show that single bacterium contact matrices from the two time points are highly similar, demonstrating the stability of both the observed 3D structures and the overall bacterial community (Fig. 3d and Supplementary Fig. 5). Although *M. instestinale* is less abundant in the second sample, the peculiar structures of its genome are still detected.

**Figure 4:**
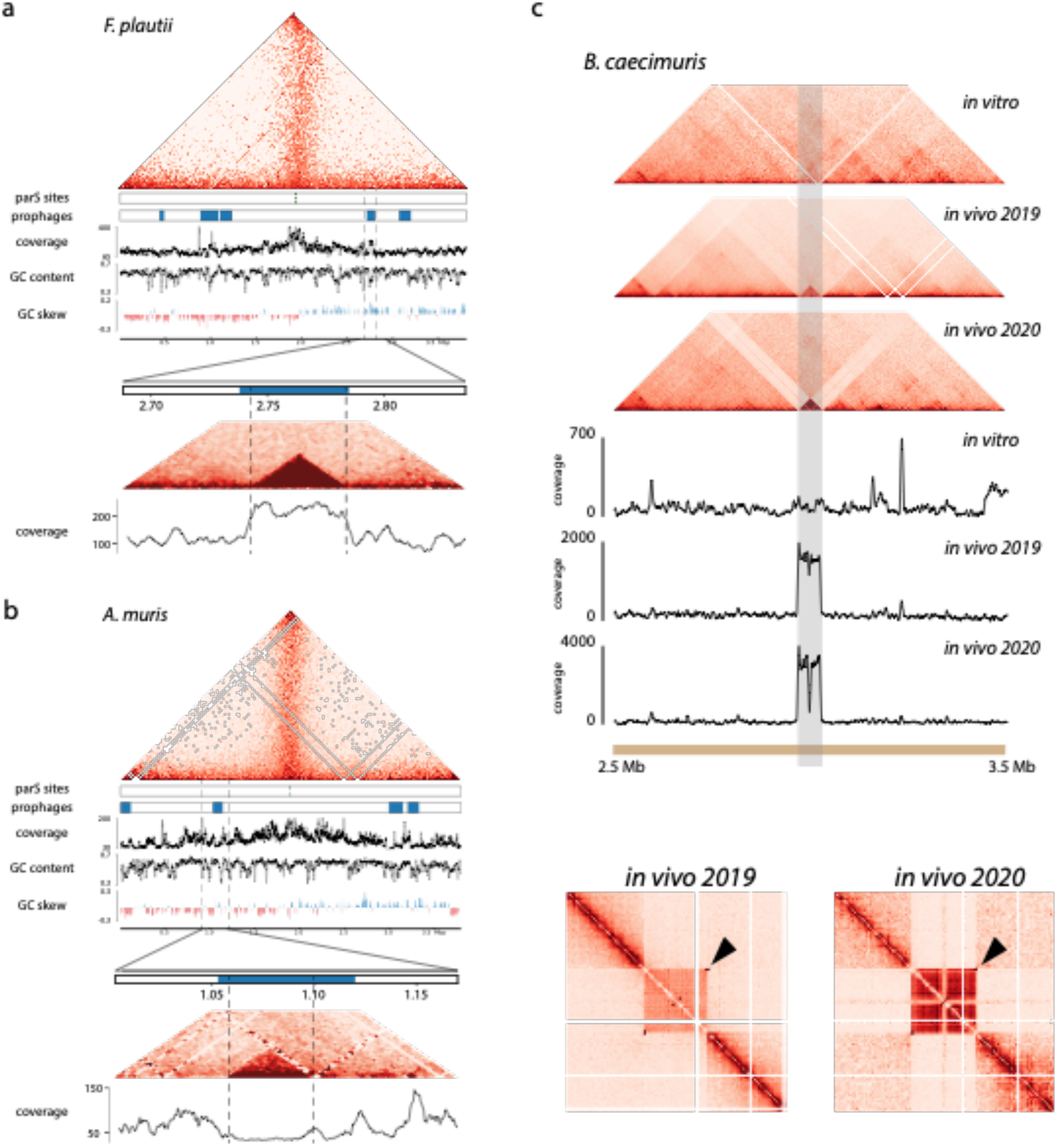
Contact maps and prophage regions in OMM^12^ bacteria. **a. b**. Contact map of *F. plautii* and *A. muris*. Contact map is shown (5 kb resolution) on top, while additional information (localization of *parS* sites, putative prophage regions, coverage, GC content, GC skew and genomic coordinates) are shown on the bottom. A zoom of the contact maps is visible on one putative prophage regions per bacteria (2 kb resolution), with the associated coverage. Prophages are visible as red squares. Hi-C signal prediction is indicated in dashed lines **c**. (upper part) Contact matrices (5 kb resolution) and associated coverage of *B. caecimuris* for *in vitro* and *in vivo* Hi-C matrices. The region corresponding to the induced prophages is highlighted in grey. (bottom part) Zoomed contact matrix, centered around the induced prophage of *B. caecimuris* (2 kb resolution). The circularization signal is highlighted by a black arrow. The depleted region of the phage is visible in the middle of the square formed by the prophage.

**Figure 5:**
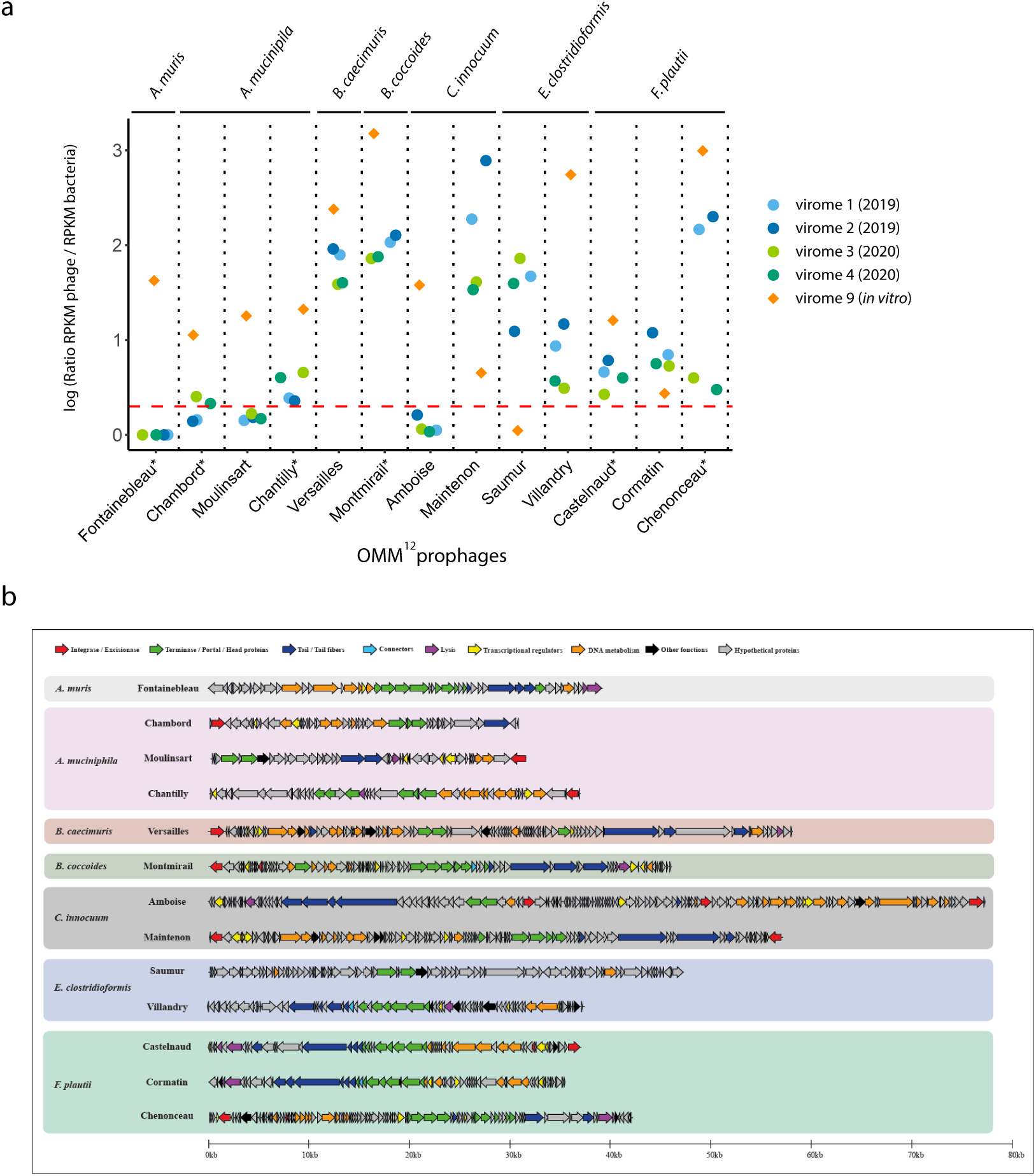
active prophages of the OMM^12^ mice. **a**. RPKM counts for the induced OMM^12^ prophages, for the different virome samples (Methods). Virome sequencing was performed for two fecal pellets of two mice in 2019 and 2020. The reads obtained were mapped on the 12 OMM^12^ strains, and the resulting normalized sequencing depth is represented. Stars aside prophages name under the graph indicate phages described by Zünd *et al*. The dashed red line indicate the induction threshold. **b**. Genetic map of the 13 induced prophages. Prophages were grouped by host. Genes were colored based on predicted functions (Methods).

Differences in the 3D organization of genomes between *in vitro* and *in vivo* conditions highlight the impact of intestinal tract on bacteria metabolism. Coupled to the stability of structures detected *in vivo*, it demonstrates that OMM^12^ mice can be further exploited to investigate chromosome architecture and dynamics in the gut environment.

### The 3D signatures of predicted prophage regions of OMM^12^ bacteria distinguish functional from cryptic prophages

Analysis of the 12 bacterial genomes using VirSorter2 (Guo et al., 2021) and VIBRANT (Kieft et al., 2020) revealed that no prophages were identified in *M. intestinale* and *B. animalis*, whereas a total of 44 prophages were predicted in the other 10 genomes, with 13 prophages in *E. clostridioformis* alone. Of these 44 candidate prophages, 14 exhibit a 3D pattern in the isolated cultures, allowing their coordinates to be refined (Table 1, Fig. 4a-b and Supplementary Fig. 6). Interestingly, 4 prophages show coverage above the median coverage of the corresponding bacterial genome, which may indicate that they are induced (YL44-pp-0.038, YL44-pp-0.712, YL32-pp-3.355 and YL31-pp-2.738) (Table 1). We noticed that the *A. muciniphila* prophage (YL44-pp-0.038) has a typical profile of duplicated sequences that correspond to strong interactions with several loci, visible as stripes in the contact matrix, which could explain the difference of coverage. Such a pattern could be the results of phage propagation at different loci in the bacterial population. For the remaining 10 prophages, the absence of a clear differential coverage associated with a 3D pattern could be the result of basal and non-abundant induction of the corresponding prophages. Interestingly, all 6 mytomicin C-induced prophages characterized previously by Zünd *et al*. (Zünd et al., 2021) exhibit a clear 3D motif. These results highlight the benefit of using Hi-C to characterize prophages, refine their boundaries and discriminate between functional and cryptic prophages. Other candidate prophages were not associated with particular 3D signatures potentially reflecting their non functionality.

We next analyzed the *in vivo* Hi-C data. Among the 14 prophages listed above, six belong to bacteria without sufficient coverage and four exhibited a similar pattern between the two conditions. In contrast, four present significant increase in coverage and/or changes in pattern compared to *in vitro* contact maps (I48-pp-2.969, YL32-pp-2.059, YL32-pp-3.355 and I46-pp-4.275) (Table 1 and Supplementary Fig. 6). The prophage I48-pp-2.969 of *B. caecimuris* presents a strong increase in its coverage associated with a clear isolation from the rest of the genome as well as a circularization signal (Fig. 4c) (Marbouty et al., 2015, 2021). These observations indicate a strong activation of this prophage. Interestingly, we observe a sharp decrease of coverage in the middle of the prophage sequence, which could be due to mapping issues or suggest that the region is deleted when the phage is induced. In *E. clostridioformis*, the prophage YL32-pp-2.059 shows an increase in coverage associated with the emergence of a circularization signal, while prophage YL32-pp-3.355 exhibits the opposite pattern. This result suggests that the gut environment switches the activation of these two prophages. Finally, prophage I46-pp-4.275 of *C. innocuum* displays a puzzling profile. Indeed, while only the left part of the prophage was observable *in vitro* as a topological domain, its right part appears as a domain under *in vivo* conditions. This suggests the presence of two functional prophages in this genomic region with different behavior depending on the environment. We also detected exclusively under *in vivo* conditions a prophage locus of *B. caecimuris* with a loop motif associated with the formation of a self-interacting domain (Table 1 and Supplementary Fig. 6). Our analysis showed that in seven bacteria, a total of 16 regions annotated as prophages exhibit 3D signatures potentially revealing functional prophages and uncovered differential prophages behavior between *in vitro* and *in vivo* conditions.

### Comprehensive characterization of the virome of the OMM^12^ mice

In order to evaluate to which extent the 3D patterns observed correlate with the production of phage particles, we performed virome sequencing (Methods) on the supernatants of *in vitro* and *in vivo* samples originating from the same cultures, same cages and at the same time as the samples used for Hi-C (Supplementary Table 1). The reads obtained were mapped on the OMM^12^ genomes and the resulting read per kilobase per million (RPKM) counts showed that multiple regions in the genomes of several strains contained significantly enriched counts, indicating that those regions are found in virus-like particles (VLPs), and thus, presumably, correspond to induced prophages (Fig. 5a). Our data showed that a total of 13 prophages from 7 strains were induced either *in vitro* (n=12) and/or *in vivo* (n=10). The six prophages previously characterized by Zünd *et al* (Zünd et al., 2021) were found among the induced prophages. Of the 16 prophages regions defined by Hi-C (15 *in vitro* and / or *in vivo* + 1 *in vivo only*), 11 were found to be among the 13 induced prophages. For the remaining 5 functional Hi-C-defined candidate prophages, we were unable to detect any significant signal in the virome data suggesting that those prophages are not active enough, or at all, in the test conditions, or are not functional or produce abortive cycles. Two induced prophages were also not found to be associated with a specific 3D pattern, possibly due to a lack of structuration of the corresponding phages in their insertion site or less likely to differences between fecal samples recovered for Hi-C and virome. We then compared the prophages coordinates defined by Hi-C data with variations of virome sequences and found that the boundaries of the prophages often agree between the datasets (9/11) (Table 2). The comparison of the virome signal from the *in vivo* samples showed minimal differences between the two mice at each time point. Importantly, the same 10 prophages were found to be induced in all four *in vivo* samples, demonstrating that the virome of the OMM^12^ mice is particularly stable between individuals and over time. While most of the prophages were found induced both *in vitro* and *in vivo*, three prophages were induced exclusively *in vitro* and one exclusively *in vivo*. Moreover, using Hi-C approach, we detected the switch of *E. colistridioformis* and *C. innocuum* prophages activation, which was confirmed by virome sequencing.

**Table 2:**
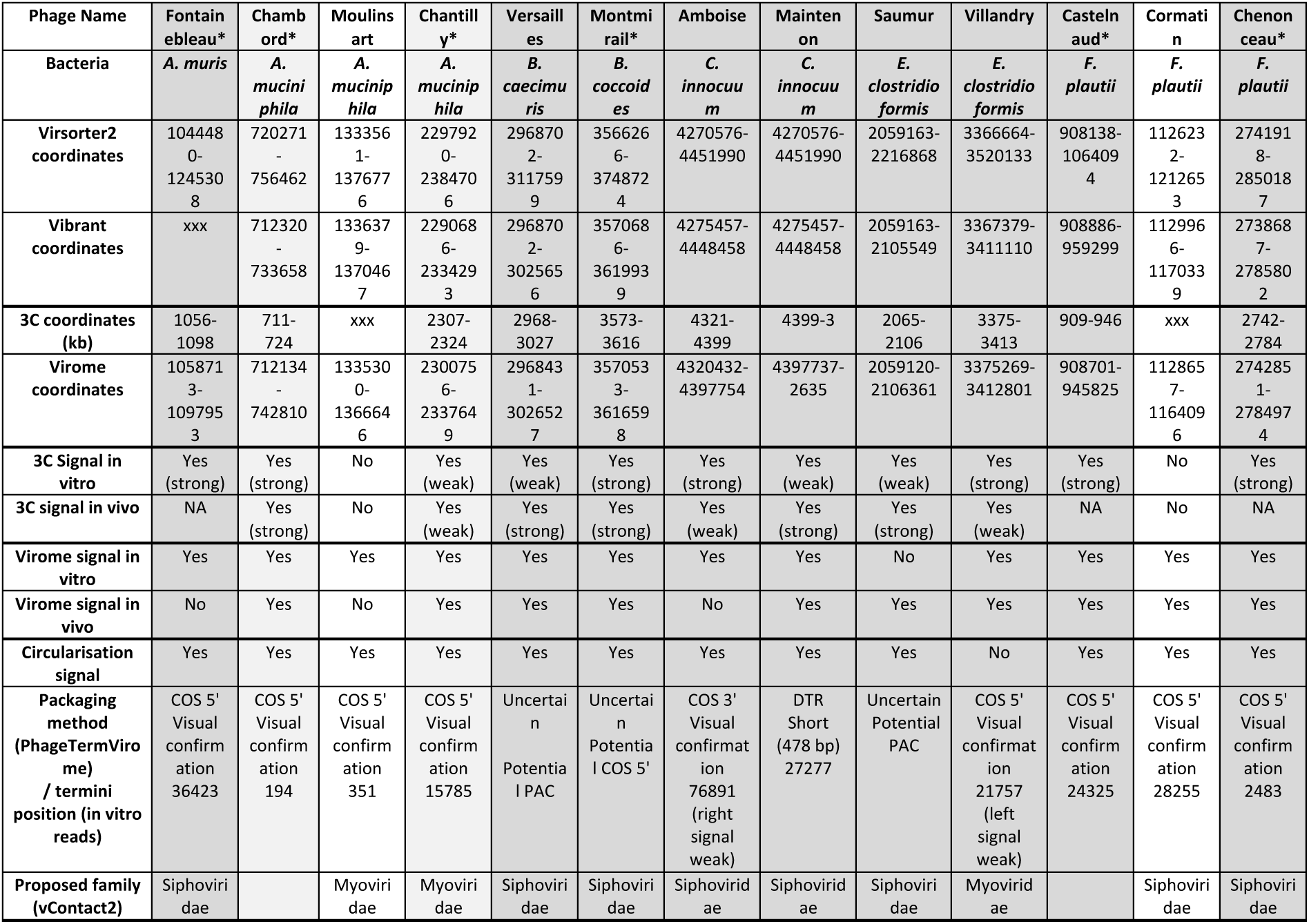
Summary of the 13 induced prophages in the OMM^12^ consortium. Grey columns indicate candidate prophages detected by Hi-C (dark grey = agreement between Hi-C and virome prophage coordinates, light grey = disagreement between Hi-C and virome prophage coordinates). Stars aside prophages name under the graph indicate phages described by Zünd *et al*.

To further characterize the virome of OMM^12^ mice, we first sequenced additional samples spiked with known concentrations of phages (Shkoporov et al., 2018). This led us to estimate that OMM^12^ fecal samples contain around 10^8^ VLP/g of feces (Methods). Second, we searched for free viruses from reads that did not match OMM^12^ genomes to perform assemblies with MEGAHIT (Li et al., 2015) and SPAdes (Antipov et al., 2020) (∼2-4% of the reads). However, this resulted in only five contigs of a size superior to 5 kb (Supplementary Table 2). BLAST analysis of these contigs revealed that they correspond to a portion of induced prophages of *B. caecimuris* or *C. innocuum*, and thus likely result from alignment errors (Supplementary Table 3). The unmapped reads were also used for taxonomic annotation using Kaiju (Menzel et al., 2016) (Supplementary Fig. 7). The vast majority of reads (99.8%) were annotated as OMM^12^ community bacteria. The remaining 0.2% of reads annotated as viral were scattered among a variety of viral phyla, showing that no particular virus was identified. Third, we explored the presence of ssDNA phages (Roux et al., 2016) (Methods), known to be abundant in the mammalian gut virome (Roux et al., 2019), but found none. Therefore, the OMM^12^ gut harbors a stable population of induced prophages and neither virulent phages nor eukaryotic viruses, providing opportunities for *in vivo* studies of exogenous intestinal viruses or endogenous murine provirus dynamics.

Altogether, our data demonstrates the ability of Hi-C as a new tool to distinguish between functional and cryptic prophages and to refine the annotation of the phage coordinates provided by state-of-the-art prediction tools. It also shows that the OMM^12^ mice represent a suitable model to study the dynamics of intestinal prokaryotic and eukaryotic viruses.

### In-depth analysis of induced prophages

The above 13 induced prophages were named after French castles and their genomes were analyzed in detail (Fig. 5b and Table 2). First, we looked for a circularization signal, in the virome sample reads (Methods). We detected such a signal for 12 out of 13 prophages, which allowed us to precisely map the prophage coordinates. Second, using those exact coordinates, we annotated the genomes using the PHROG database (Terzian et al., 2021) (Methods). Between 25% to 60% of the genes were annotated, and we identified terminase/portal proteins, two hallmarks of phage proteins, in all genomes (Fig. 5b). Integrases were also frequently found, in agreement to the temperate lifestyle of these phages. Third, PhageTermVirome (Garneau et al., 2021) was successfully used for 10 phages to confirm the viral nature of these prophages as well as to identify their packaging method and termini. (Table 2). Finally, we searched for homologs using BLAST, but only distant homologs were identified, showing that these prophages are relatively new. Thus, we used vContact2 (Bin Jang et al., 2019) to generate shared protein networks and identify the groups of phages that most closely relate to these 13 phages. While only few high-quality RefSeq genomes clustered closely to the OMM^12^ prophages, many metagenomic phage sequences belonging to the Cenote Human Virome Database (CHVD) (Tisza and Buck, 2021) closely clustered to them (Supplementary Figure 8). This clustering allowed us to predict their possible morphology (Table 2). Overall, these 13 induced OMM^12^ phages represent newly characterized viruses from representative of intestinal bacteria, from which the assignation of a number of viral reads from metagenomics studies will contribute to lower the “viral dark matter”.

## DISCUSSION

In the present study, we combined Hi-C and virome sequencing to dissect the chromosome architecture of a synthetic bacterial community representative of the murine gut microbiota and identify its associated induced prophages in both *in vitro* and *in vivo* conditions. Most of the Hi-C studies have so far focused on model bacteria encompassing mainly representative of the proteo- and actinobacteria and were performed exclusively on *in vitro* grown cultures (Böhm et al., 2020; Lioy et al., 2020; Szafran et al., 2021; Tran et al., 2017). The Hi-C results from the 12 non-model bacteria that belong to major phylum of the mammals gut confirmed the conservation of the ParABS system and the condensins in the organization and the dynamics of bacterial chromosome architectures. On the other hand, the exact role of the Smc-ScpAB complex remains puzzling as some species encoding for this complex do not display a strong opposite diagonal (*L. reuteri* and *E. faecalis*) in their contact maps. It suggests that this complex may not be functional or is involved in a novel way that does not necessarily lead to an alignment of the chromosomal arms. Moreover, we also detect for *B. caecimuris* and *T. muris* a weak signal in the opposite diagonal while no clear homologs of condensins could be characterized in their genomes. As it is thought that condensins are essential actors of chromosomes organization (Mäkelä and Sherratt, 2020), our results suggest that other condensins related proteins remain to be uncovered in bacteria. Taken together, we uncovered a diversity of architecture and genomic organization by studying non-model bacteria and linking 3D organization with physiological state (*in vitro* vs *in vivo* growth).

We also found that the global chromosome folding of the 12 bacteria are preserved in the gut compared to *in vitro* condition with, still, notable differences at the local scale. Some structures appear less pronounced in the contact matrices obtained with fecal samples and the overall ratio of short- and long-range interactions appear to change between the two conditions. This result is likely due to differences in the growth conditons imposed by the gut environment and demonstrates its high impact on bacterial physiology. Such observation is in line with contact maps obtained from stationary phase cultures that exhibit less pronounced signal in the main diagonal but higher short-range interactions compared to exponential growth condition (Lioy et al., 2018). However, this time the ratio of the ori/ter coverage indicates that bacteria are dividing. Future studies combining Hi-C with RNAseq will provide a better understanding of the link between chromosome 3D architecture, transcriptional activity and the physiological state of the bacteria in the intestinal tract. Given the stability over time of the observed structures, the OMM^12^ mice represent a very promising model to study the genome dynamics of intestinal microbial communities at a very high resolution.

Prophages activity in host bacteria are generally studied either by qPCR or by exploiting sequencing reads from enriched virions or fecal samples and using phages reference genomes (Zünd et al., 2021). We previously found that the Hi-C data could be exploited to study prophage in bacteria (Marbouty et al., 2015, 2017). Here, we leverage Hi-C with virome data to detect a specific 3D pattern for 16 prophages among the 44 candidates predicted from genome analysis. Among these 16 prophages, we found that 11 were indeed induced *in vitro* and/or *in vivo*. We also found that only two induced prophages did not display a specific 3D pattern. Of particular interest is the use of the Hi-C data to refine the borders of the prophage loci predicted by state-of-the-art tools. Moreover, our approach also offers the possibility to detect circular or loop signal that can help detecting prophages activation and could as well allow the characterization of their replication and packaging strategy.

We found seven additional phages induced *in vitro* and / or *in vivo* compared to the six mitomycin C-induced prophages previously reported from individual cultures (Zünd et al., 2021). This shows that the SOS response induced by DNA damages is not an universal way of inducing prophages (Cornuault et al., 2020). In addition, some prophages were differential induced between the two conditions, with three only induced *in vitro* and one only induced *in vivo*. These differences could be driven by the gut environment or the contact with the other bacteria. Previous studies have documented particular prophage dynamics in the gut of mice (Cornuault et al., 2020; De Paepe et al., 2016; Oh et al., 2019) but the impact of the intestinal environment on prophage induction remains poorly understood. Here, we showed that the resident viral community of the OMM^12^ mice is very stable over time, being uniquely constituted of temperate phages, which is consistent with the high stability of the bacteriome in the these mice (Eberl et al., 2020). This is also coherent with the current appreciation of the human virome, which is thought to have a minimal intra-personal variability (Shkoporov et al., 2019). These results advocate for the OMM^12^ model to study of intestinal viruses dynamics.

This first characterization of the viral community of gnotobiotic animals showed that the OMM^12^ mice do not carry any eukaryotic intestinal virus. Such viruses are part of the usual mouse microbiota (Rasmussen et al., 2021; Schulfer et al., 2020), and have been shown to contribute to the development of immunity (Dallari et al., 2021; Ingle et al., 2019). Their absence might have consequences on the maturation of the immune system but also offers opportunities to study this process for individual viruses (pathogens or commensals) in the presence of a defined intestinal microbiota. In addition, we did not detect virulent phages in the virome samples. The absence of virulent phages and eukaryotic viruses are however not surprising considering the stringent conditions in which these mice are bred.

Altogether, these data, obtained with an gnotobiotic murine model, have demonstrated the benefit of the Hi-C approach to not only unveils novel chromosome architectures but also to detect the dynamic induction of prophages. Coupled to virome analysis, we performed the most extensive characterization of the dynamic variations of the bacteriome and the virome of intestinal communities, which provide a path to move microbiota studies from correlation to causality.

## ACKNOWLEDGEMENTS

We thank Cyril Matthey Doret and Agnès Thierry for assistance during computational and experimental work, respectively. We also thank Marie-Agnès Petit for her precious advices and help in prophages annotation. We thank Lorenzo Chaffringeon for fruitful discussions. This research was supported by funding to BS from DFG-STE-1971/11-1 (PhaStGut project); to BS from the European Research Council under the Horizon 2020 Program (ERC grant agreement 865615), to LD and MMa from PRCI ANR-20-CE92-0048 (PhaStGut project); to RK from the European Research Council under the Horizon 2020 Program (ERC grant agreement 771813) and from JPI-EC-AMR STARCS ANR-16-JPEC-0003-05. QLB. received funding by École Doctorale FIRE-Program Bettencourt. AB is supported by an ENS fellowship by the French Ministry of Higher Education, Research and Innovation. MMo and JRG were supported by JCJC ANR-18-CE35-0011 (project CDPhages). MT received funding from DigestScience. Biomics Platform, C2RT, Institut Pasteur, Paris, France, was supported by France Génomique (ANR-10-INBS-09) and IBISA.

## DATA AND CODE AVAILABILITY

Sequence data as well as new bacterial genomes have been deposited in the NCBI under the BioProject number PRJNAXXX.

Phages genomes are available using the following project number: PRJNAXXX.

Code used in the present study can be found at the following address https://github.com/

## ETHICS STATEMENT

A total of 8 OMM^12^ mice (3 females; 5 males; seven to nine weeks old) reared at Institut Pasteur (Paris, France) and housed in the gnotobiotic facility in accordance with Institut Pasteur guidelines and European recommendations. Food and drinking water were provided ad libitum. Protocols were approved by the veterinary staff of the Institut Pasteur animal facility (Ref.#18.271) and the National Ethics Committee (APAFIS#26874-2020081309052574 v1).

## AUTHORS CONTRIBUTIONS

Conceptualization: MMa, LD, RK, BS; Experiments: QLB, MMa, DC, MT, AvS, JRG; informatic analysis: QLB, AB, MMa, JRG; funding acquisition: MMa, LD, RK, BS, MMo; Writing: MM, LD, QLB, RK, DC, JRG, AB, BS.

## MATERIAL AND METHODS

### Bacterial culture

The OMM^12^ strains are originating from the miBC collection (Lagkouvardos et al., 2016). All bacteria were grown in anaerobic akkermansia medium (18,5 g.L^-1^ brain heart infusion, 15 g.L^- 1^ trypticase soy broth, 5 g.L^-1^ yeast extract, 2.5 g.L^-1^ K_2_HPO_4_, 0.5 g.L^-1^ cysteine hydrochloride, 0.5 g.L^-1^ glucose, 0.4 g.L^-1^ Na_2_CO_3_, 1 mg.L^-1^ hemin, 0.5 mg.L^-1^ menadione and 3% fetal calf serum, completement-inactivated, in distilled water) in an anaerobic chamber (1.5 - 3 % H_2_, 4% CO_2_, rest N_2_). 60 mL of medium were inoculated to reach a OD_600nm_ of 0.01 for each bacterium and were incubated without agitation at 37°C. *Muribaculum intestinale* (YL27), *Clostridium innocuum* (I46), *Blautia coccoides* (YL58), *Limosilactobacillus reuteri* (I49), *Enterococcus faecalis* (KB1) and *Bifidobacterium longum subsp. animalis* (YL2) were cultured for 5 h, while *Bacteroides caecimuris* (I48), *Akkermansia muciniphila* (YL44), *Enterocloster clostridioformis* (YL32), *Flavonifractor plautii* (YL31), *Turicimonas muris* (YL45) and *Acutalibacter muris* (KB18) were cultured for 10 h. Half of each culture was centrifuged (6,000 g, 15 min, 4°C). The supernatant was frozen at -80°C. Formaldehyde was added to the other half (3% final) and the mixture was incubated under gentle agitation at room temperature for 30 min, then at 4°C for 30 min. 5 mL of glycine 2.5 M was then added, followed by an incubation at room temperature for 20 min under gentle agitation. The solution was centrifuged (6,000 g, 10 min, 4°C) and the pellet was washed in 1X PBS. After a similar centrifugation, the supernatant was removed, and the pellet was frozen at -80°C.

### Hi-C libraries generation

Hi-C libraries were generated as previously described (Marbouty et al., 2021). Mice fecal samples were collected and directly mixed in 10 mL of crosslinking solution (1X PBS supplemented with 3% formaldehyde) and incubated for one hour at room temperature under strong agitation. Formaldehyde was quenched by adding 5 mL of 2.5 M glycine during 20 min at room temperature under gentle agitation. Samples were then recovered by centrifugation (6,000 g, 10 min, 4°C), washed with 10 mL 1X PBS, re-centrifuged and stored at −80°C until processing. Each sample was resuspended in 1.2 mL TE 1X supplemented with antiprotease (mini tablets - Roche), transferred in a Precelys tubes (2 mL – VK05 supplemented with 100 µL of VK01 glass beads) and disrupted (6,700 rpm – 20s ON / 30s OFF – 6 cycles). Lysates were recovered and SDS 10% was added to a final concentration of 0.5% and the lysate was incubated 10 min at RT. For each library, 1 mL of lysate was transferred to a tube containing the digestion reaction solution (500 µL NEB1 10X buffer, 500 µL Triton 10%, 1000 U Sau3AI, H_2_O, final volume = 4 mL). Digestion was allowed to proceed for 3 h at 37°C under gentle agitation. Tubes were then centrifuged for 20 min at 4°C and 16,000 g and the supernatants discarded and pellets were resuspended in 400 µL H2O. Biotinylation was done by adding 50 µL NEB ligation buffer 10X (without ATP), 4.5 µL of 10 mM dATP/dTTP/dGTP, 37.5 µL biotin-dCTP 0.4 mM and 8 µL Klenow (5U / µL). Reactions were incubated for 45 min at 37°C and then transferred to a tube containing the ligation reaction (160 µL NEB ligation buffer 10X, 16 µL ATP 100 mM, 16 µL BSA 10 mg/mL, 500 U T4 DNA ligase, final volume = 1.1 mL). Ligations were processed for 3 h at RT. 20 µL EDTA 0.5M, 80 µL SDS 10% and 2 mg proteinase K were added to each reaction and incubated overnight at 65°C to digest proteins. DNA was extracted using phenol-chloroform and precipitated with 2.5 x vol. ethanol 100%. Pellets were suspended in a final volume of 130 µL TE 1X supplemented with RNAse, incubated 1 h at 37°C and stored at -20°C until use. DNA was extracted, purified and processed into sequencing library as described previously (Moreau et al., 2018). Proximity ligation libraries were sequenced using pair-end (PE) Illumina sequencing (2 × 35 bp, NextSeq500 apparatus) (Supplementary Table 1).

### Hi-C analysis

Forward and reverse reads were aligned separately with bowtie2 v2.3.5.1 (Langmead and Salzberg, 2012). From these alignments, Hi-C matrices and genomic distance law were generated using hicstuff v3.0.3 (https://github.com/koszullab/hicstuff). Matrices were balanced using ICE algorithm (Imakaev et al., 2012). For comparative analysis, matrices were binned at 10 kbp resolution and were downsampled to the same number of contacts. Reproducibility between replicates was assessed using the hicreppy v0.0.6 implementation (https://github.com/cmdoret/hicreppy) of the HiCrep algorithm (Yang et al., 2017). Comparison between matrices were done using log2 ratio and serpentine v0.1.3 60 for flexible binning (Baudry et al., 2020).

### Bacterial genomes annotations and genomic features

Genomes were annotated using the last version of the NCBI Prokaryotic Genome Annotation Pipeline (PGAP) (Li et al., 2021). The coverage was computed using tinycov (https://github.com/cmdoret/tinycov), v0.3.0. The GC content and the GC skew were computed using dnaglider (https://github.com/cmdoret/dnaglider), v0.0.4. The *parS* sites have been detected using a degenerated *par*S consensus sequence TGTTTCACGTGAAACA (Livny et al., 2007) and allowing 2 mismatches. The ori and ter position were approximated based on the GC skew inversion. The shift closer to the *par*S cluster were annotated as the ori and the shift as the opposite of the genome as the ter. Figures were generated using pyGenometracks v3.6 (Lopez-Delisle et al., 2021).

### Prophage annotations

Prophage annotation was performed using the OMM^12^ genomes (Lamy-Besnier et al., 2021) except for *B. caecimuris, B. longum* and *F. plautii* for which the version re-assembled in this article was used. Both Vibrant (Kieft et al., 2020), v1.2.1 and Virsorter2 (Guo et al., 2021), v2.2.3 with their respective databases, were used to annotate the bacterial genomes. The data from mitomycin C-induced prophages was obtained from Zünd *et al*. (Zünd et al., 2021).

### Virome preparation

Fecal samples (2 pellets minimum) were collected and directly frozen at -20°C. The pellets were then resuspended in 14 mL Tris 10 mM, pH 8 and centrifuged for 10 min at 5,200g, 4°C. A known number of phages were then added as spike as indicated in Supplementary Table 1. The supernatant was then filtered (0.45 µm and 0.22 µm) and ultracentrifuged for 3 h at 270,000 g. The resulting pellet was resuspended in 500 µL of TN buffer (10 mM Tris, 150 M, pH 7.5). To remove free DNA and RNA, 4 U of Turbo DNaseI (Ambion) and 10 µL of RNAse (A/T1 mix, ThermoFisher) were added for 30 min at 37°C. The DNAse was then inactivated with 15 mM final EDTA and treated with 100 µg/mL final of proteinase K (Eurobio) and 0.5% final SDS for 30 min at 55°C. The viral DNA was extracted by adding a volume of phenol-chloroform-isoamyl alcohol (25:24:1), vortexing for 30 sec and centrifuging for 5 min at 12,000 g. The aqueous phase was recovered and treated again with phenol-chloroform-isoamyl alcohol similarly. The recovered DNA precipitated with sodium acetate (300 mM final) and two volumes of 100% EtOH. 1 µL of glycogen was also used as a DNA carrier. The sample was mixed by inversion and incubated for 2 h at -80°C before centrifugation for 20 min at 15,000 g, 4°C. The pellet was dried and resuspended in 20 µL Tris 10 mM, pH 8. The DNA concentration was measured with Qubit (Invitrogen). The dsDNA libraries were prepared using TruSeq Nano DNA Sample Preparation Kit (Illumina), and ssDNA/dsDNA libraries with the Accel-NGS™ 1S Plus DNA Library Kit (Swift Biosciences). They were sequenced on a NextSeq550 or Novaseq (Illumina) for the 2 × 35 nt libraries, and on a MiSeq (Illumina) for the 2 × 150 libraries (Supplementary Table 1).

The supernatant of individual OMM^12^ strain cultures was filtered (0.45 µm and 0.22 µm) and ultracentrifuged for 3 h at 270,000 g. The following steps were similar to the treatment of fecal samples described above. The DNA obtained for each strain was mixed in a single sequencing run, on a MiSeq (Illumina) with a 2 × 150 library.

### Annotation of induced prophages

The quality of the reads was assessed with FastQC (https://www.bioinformatics.babraham.ac.uk/projects/fastqc/). The reads were cleaned with cutadapt v2.10 (Martin, 2011) using a quality threshold of 20 and a minimum size of 30 for 35 bp reads or 140 for 150 bp reads. Using all the virome libraries, the precise coordinates of the prophages were determined by visualizing individual reads on IGV (Robinson et al., 2011) v2.11.9 and looking for paired-end reads mapping both at different locations of the bacterial genome in areas bioinformatically predicted as prophage regions. The observed circularization signal indicated the production of viral particles and thus the coordinates of the phage genome. Using those coordinates, the precise sequence of each prophage was extracted and annotated as follows. A first annotation was realized using PATRIC (Davis et al., 2020), with the “phage” recipe. Then, each predicted protein was compared to the PHROG (Terzian et al., 2021) database v3 using the HHsuite v3.3.0 (hhsearch_omp function). For hits with a probability above 88%, the corresponding PHROG annotation was manually added to the corresponding protein. The annotated genomes were then visualized using Clinker (Gilchrist and Chooi, 2021) v1.1.0.

### RPKM counts of induced prophage regions in OMM^12^ virome samples

The virome reads were mapped using bowtie2 (Langmead and Salzberg, 2012) v2.3.5.1 with default parameters against the most recent version of each of the 12 reference genomes of the OMM^12^, except for *B. caecimuris, F. plautii* and *B. animalis* for which the version attached to this article was used. The number of reads mapping on each prophage was divided by the number of reads (per million) and by the length of the prophage region (per kb), resulting in RPKM counts for each sample and prophage region.

In order to estimate the number of VLP present in the virome samples, the virome reads were similarly mapped on the genome of the spiked phages (CLB_P1: KC109329, CLB_P2: OL770107, CLB_P3: OL770108, M13: NC_025824). RPKM counts were similarly calculated and compared to the concentration of phage spike in the corresponding sample (Supplementary Table 1) to obtain the estimation of the number of VLP present in OMM^12^ fecal samples. This value was then divided by the initial weight of the fecal sample used for the virome protocol to obtain VLP/g values. A value of 2.47×10^8^ PFU/g was obtained for Virome 7, and 1.09×10^8^ PFU/g for Virome 8.

### Prediction of phage termini and packaging mechanism

PhageTermVirome (PTV) (Garneau et al., 2021) v.4.0.0 was run using virome sequencing reads that mapped to each inducible prophage region. PTV was run in paired-end mode, with seed parameter -s 15 and peak merging parameter -d 8. Packaging mechanisms were predicted both by statistical analysis and visual confirmation of coverage patterns near termini.

### Shared-proteins network analysis

OMM^12^ induced prophages were clustered using vConTACT2 (Bin Jang et al., 2019) v0.9.19 with the following parameters: --rel-mode ‘Diamond’ --db ‘ProkaryoticViralRefSeq201-Merged’ --pcs-mode MCL --vcs-mode ClusterONE. Publicly available phage genomes from ViralRef Seq V.201 (Brister et al., 2015) and Cenote Human Virome Database (CHVD) (Tisza and Buck, 2021) were used as reference collections for the analysis. The resulting network was visualized and annotated using Cytoscape v3.9.1 (Kohl et al., 2011). The edge-weighted spring-embedded model was applied to position genomes sharing most protein clusters.

### Search for virulent phages in OMM^12^ fecal virome

Cleaned virome reads were mapped on both the genomes of the OMM^12^ strains and the Mus musculus genome (NC_000067.7). The non-mapping reads were then used to perform assemblies with MEGAHIT v1.2.9 (Li et al., 2015) and SPAdes v3.15.2 (Nurk et al., 2017), both with default parameters. The resulting contigs <5 kb were discarded, and the other contigs were analyzed using PATRIC (Davis et al., 2020) and BLAST (Altschul et al., 1990). The same non-mapping reads were also analyzed by Kaiju v1.7.3 (Menzel et al., 2016) in order to obtain taxonomic assignation. The results were visualized using Krona (Press et al., 2017).

### Detection of ssDNA phage in virome samples

Cleaned reads from virome 5 and virome 6 (Supplementary Table 1) were merged into one single dataset. Reads were assembled using SKESA (Souvorov et al., 2018) v2.4.0 and Metaviral SPAdes (Antipov et al., 2020) v3.15.2 using k-mer size 21 and resulting contigs (≤ 15 kb ≥ 1 kb) were analyzed with VIBRANT (Kieft et al., 2020) v1.2.1 and VirSorter2 (Guo et al., 2021) v2.2.3 with default option to predict ssDNA sequences. Except for the M13 spiked control, no ssDNA was identified in the sequenced samples. All contigs from the SKESA and Metaviral SPAdes assemblies were joined in a single multifasta file and proteins were predicted and annotated using PROKKA (Seemann, 2014). The whole set of predicted proteins was then screened with BlastP for Microviridae characteristic MCP and Rep proteins contained in the PHROG database (Terzian et al., 2021) (v3, PHROG 514 and 713 respectively). No significant hits were obtained on the database using a e-value threshold of 1×10^−3^.

