## Supplementary Figures and Tables for "Chromosome folding and prophage activation reveal gut-specific genome dynamics of bacteria in the OMM^12^ consortium"

**Supplementary Figure 1a and 1b:** Contact map of each bacterial strain of the OMM$^{12}$ consortium obtained from *in vitro* cultures.
**Supplementary Figure 2**: Re-assembly of *B. animalis*, *F.* plautii, and *B. caecimuris*.
**Supplementary Figure 3:** Signal of the secondary diagonals for the different bacteria of the OMM$^{12}$ consortium.
**Supplementary Figure 4:** Comparison of the contact maps (*in vitro* vs. *in* vivo) for the six most abundant bacteria.
**Supplementary Figure 5:** Hierarchical clustering of the different Hi-C replicates for the different bacteria of the OMM$^{12}$ consortium using the software HiCrep.
**Supplementary Figure 6a and 6b:** Contact maps of functional prophage candidates (+/- 50 kb).
**Supplementary Figure 7:** Krona representation of the Kaiju annotation of the reads not mapping on the OMM$^{12}$ strains' genomes.
**Supplementary Figure 8:** Viral clustering of the 13 induced phages using vContact2.

**Supplementary Table 1:** Genomic libraries generated.
**Supplementary Table 2:** Metrics of the assemblies obtained with virome reads that did not map on the OMM12 strains.
**Supplementary Table 3:** Blast results of the contigs obtained by assembling non-mapping reads.

**Supplementary Figure 1a and 1b: Contact map of each bacterial strain of the OMM$^{12}$ consortium obtained from *in vitro* cultures.**

Each contact map is represented with associated genomic information: localization of *parS* sites (green), tRNA (blue) and rRNA (red) on top with below prophage annotation (blue), coverage, GC content and GC skew.

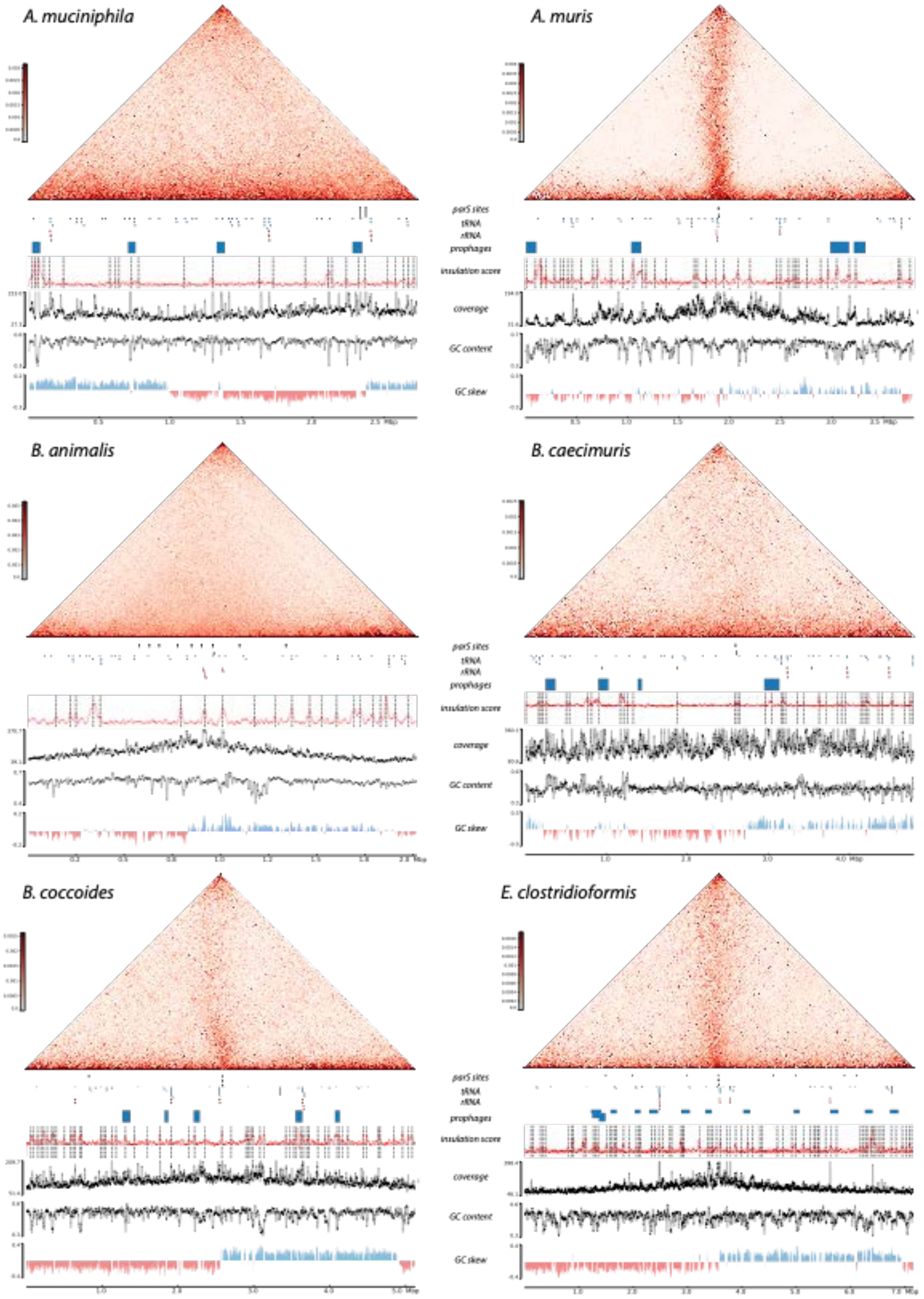

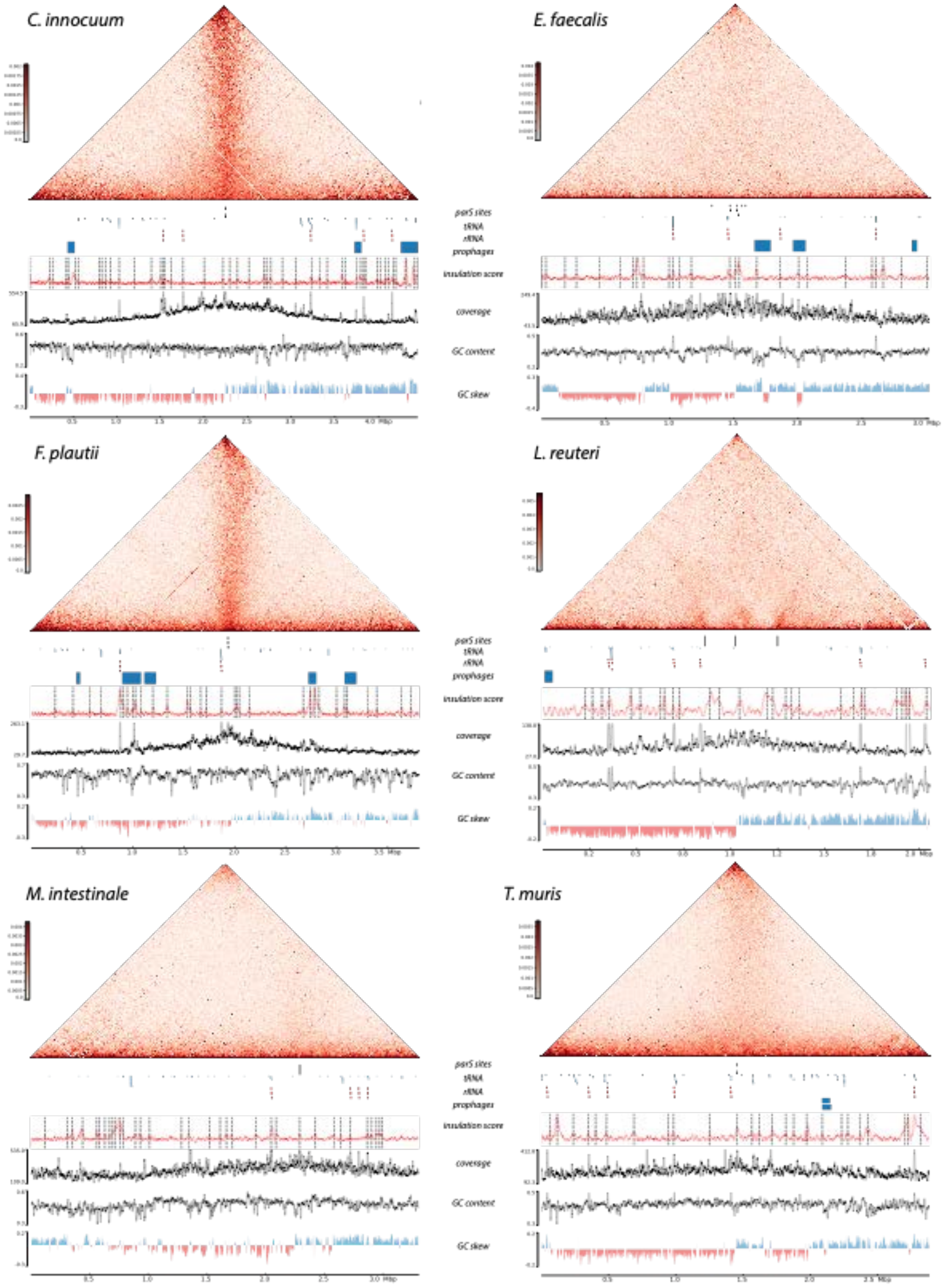
C. innocuum
E. faecalis
F. plautii
L. reuteri
M. intestinale
T. muris
parS sites
tRNA
rRNA
prophages
insulation score
coverage
GC content
GC skew
Mbp

**Supplementary Figure 2: Re-assembly of *B. animalis*, *F.* plautii, and *B. caecimuris*.**
The contact maps are shown before (left) and after (right) Hi-C-based re-assembly.

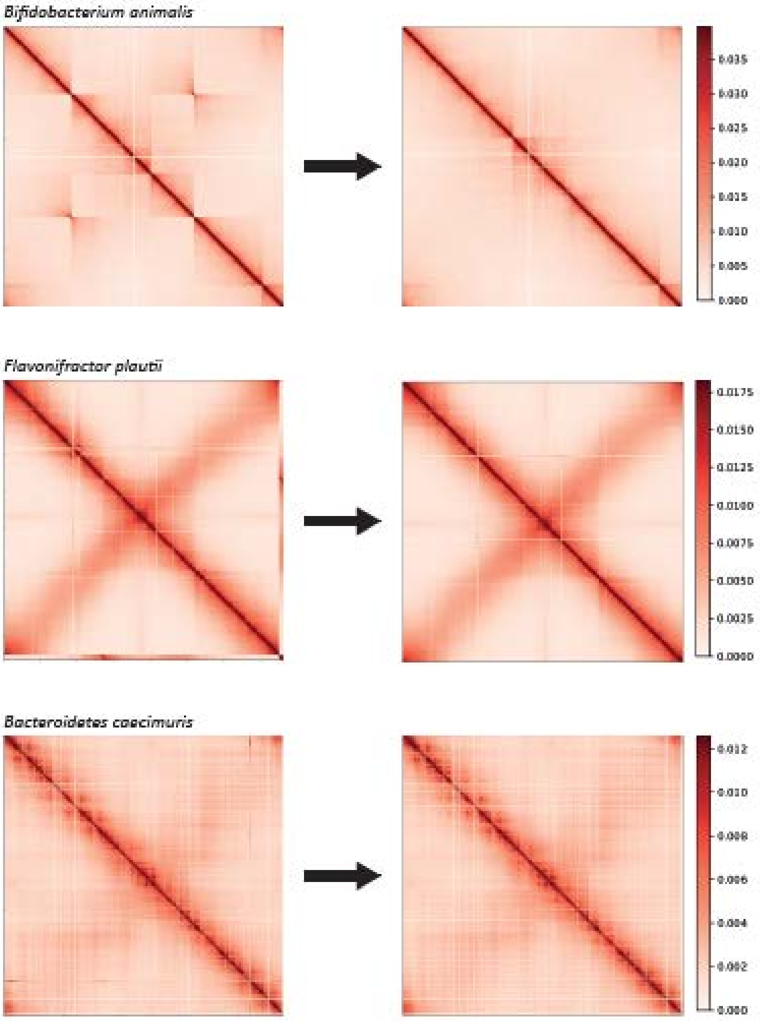

**Supplementary Figure 3: Signal of the secondary diagonals for the twelve bacteria of the OMM$^{12}$ consortium grown *in vitro*.**

Plots are centered on *ori*. Localisation of *parS* sites are indicated as red dashed lines. Red stars indicate the presence of several *parS* sites in the same 5 kb window.

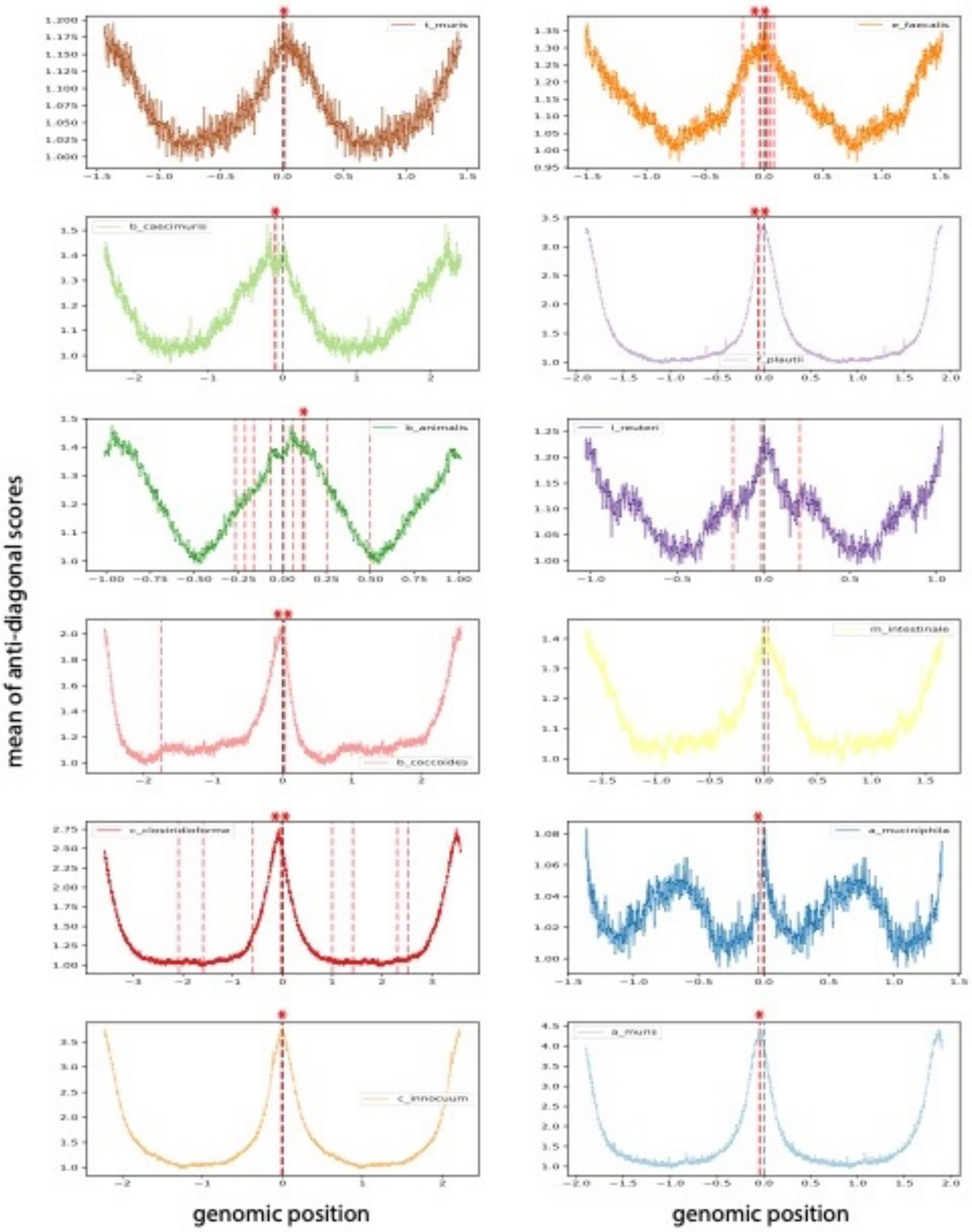

**Supplementary Figure 4: Comparison of the contact maps (*in vitro* vs. *in vivo*) for the six most abundant intestinal bacteria in OMM[12] mice.**

**a.** *In vitro* (5 kb bin), *in vivo* (5 kb bin) and ratio (Log2; 10 kb bin) of contact maps (*in vitro vs. in vivo*) obtained for the six most abundant bacteria in OMM[12] mice. Specific annotations are indicated under matrices. Origin of replication are indicated by dashed black lines. **b.** The three matrices (*in vitro* (up), *in vivo* 2019 (middle) and *in vivo* 2020 (bottom)) are shown on top of each other.

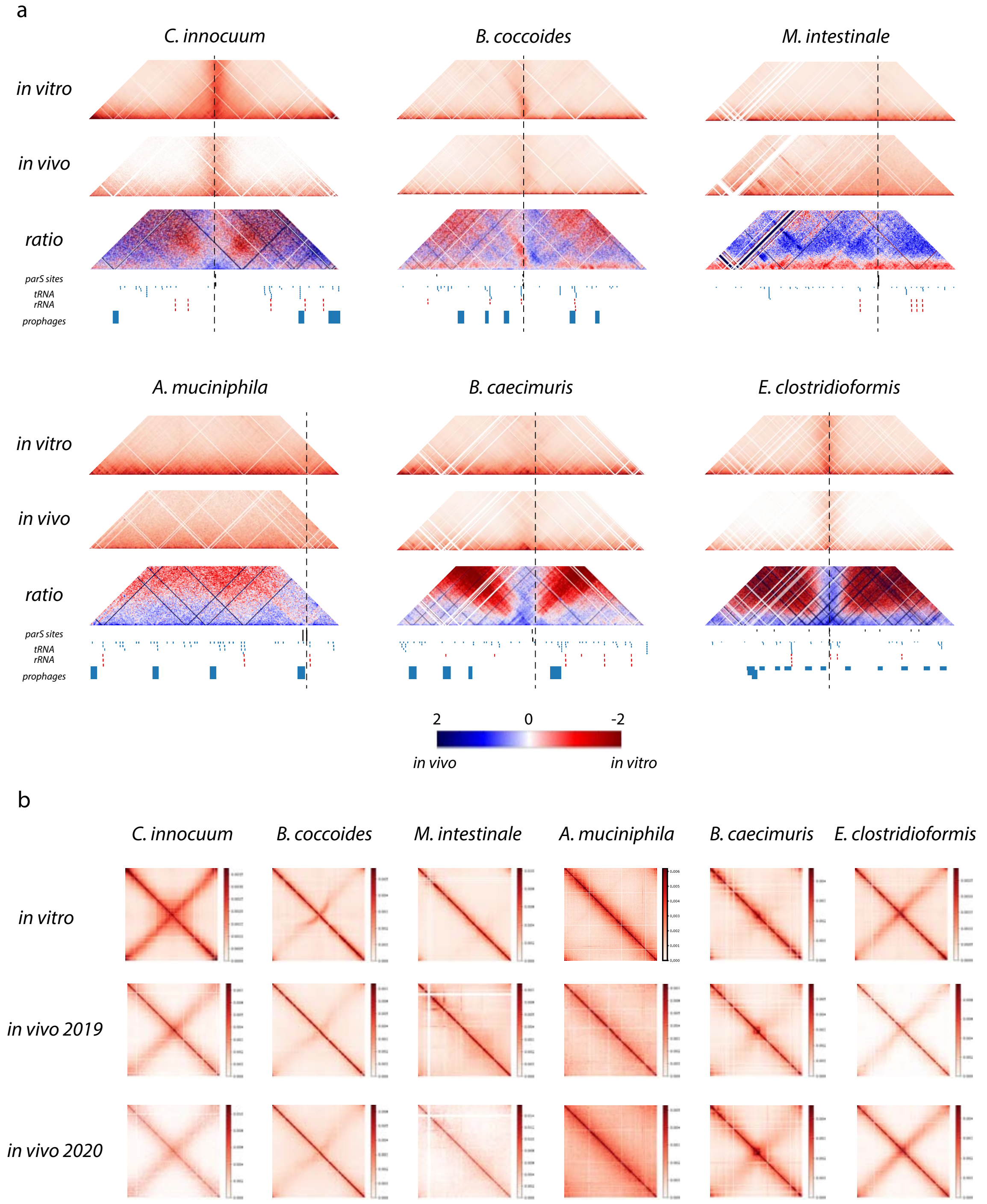

**Supplementary Figure 5: Hierarchical clustering of the different Hi-C replicates for the different bacteria of the OMM$^{12}$ consortium using the software HiCrep.**

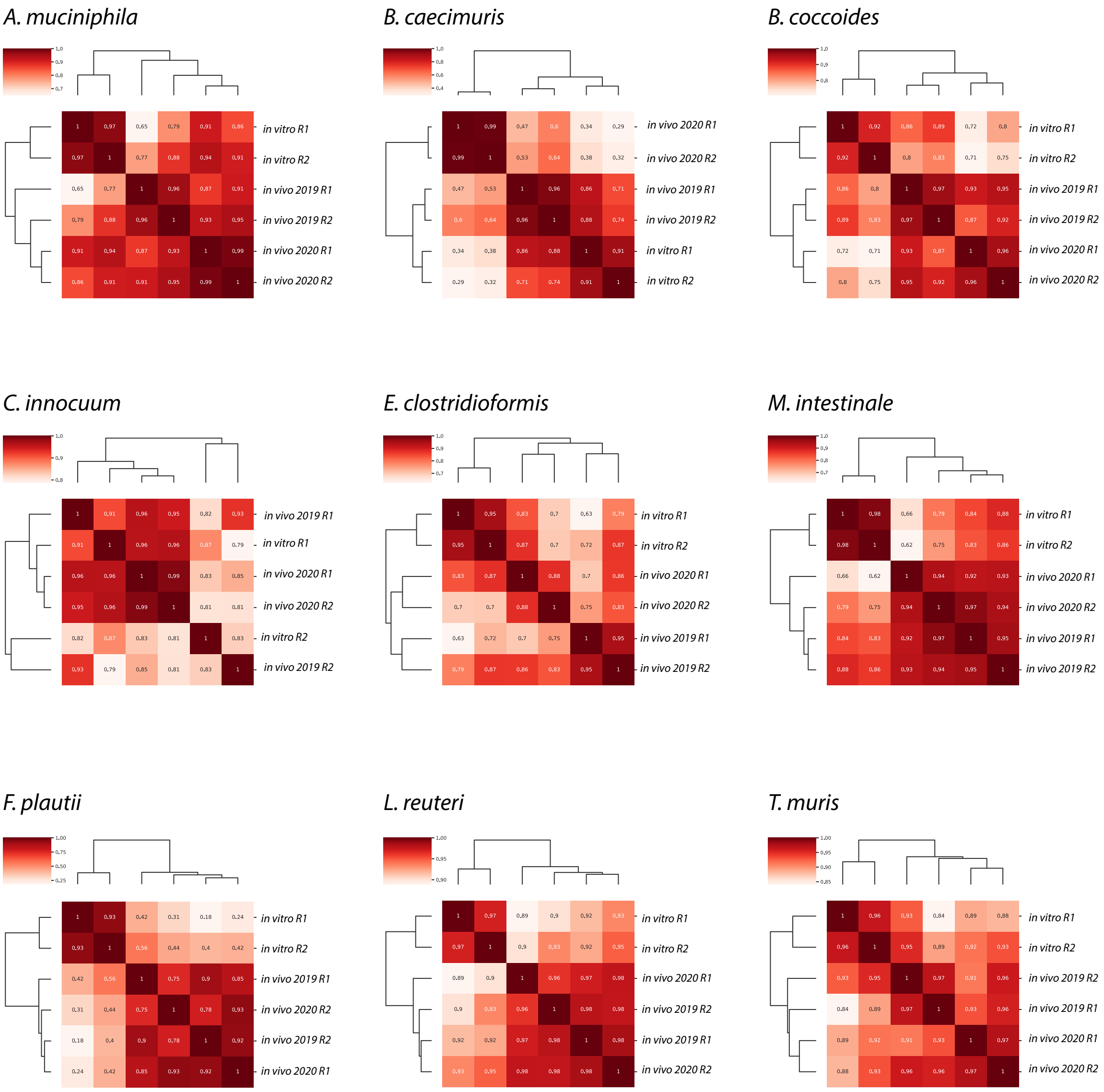

**Supplementary Figure 6a and 6b: Contact maps of functional prophage candidates (+/- 50 kb).**
Genomic coordinates are indicated above contact maps while Hi-C coverage as well as virome data are indicated under. Dashed lines indicate Hi-C refinement of prophage coordinates.

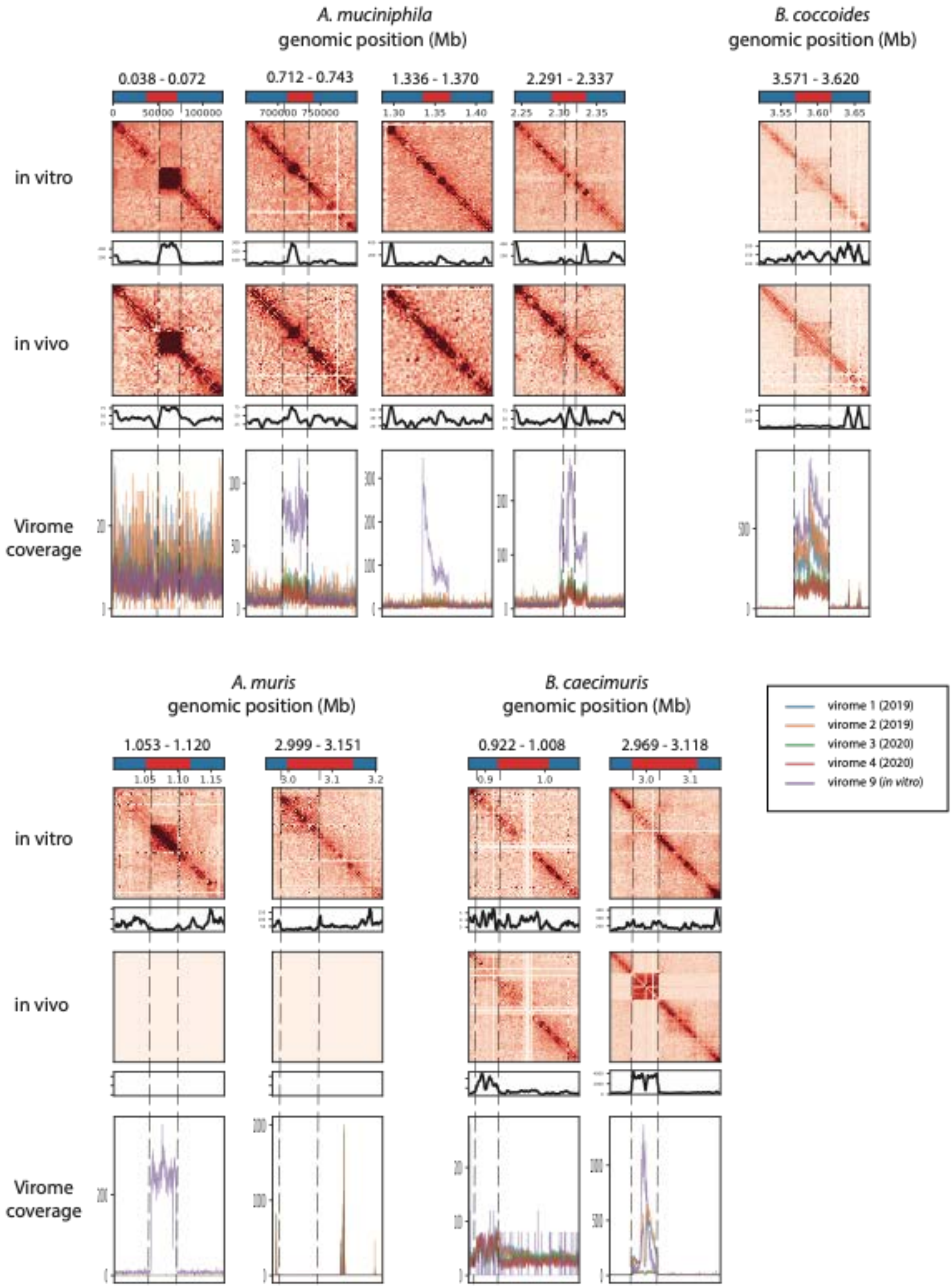

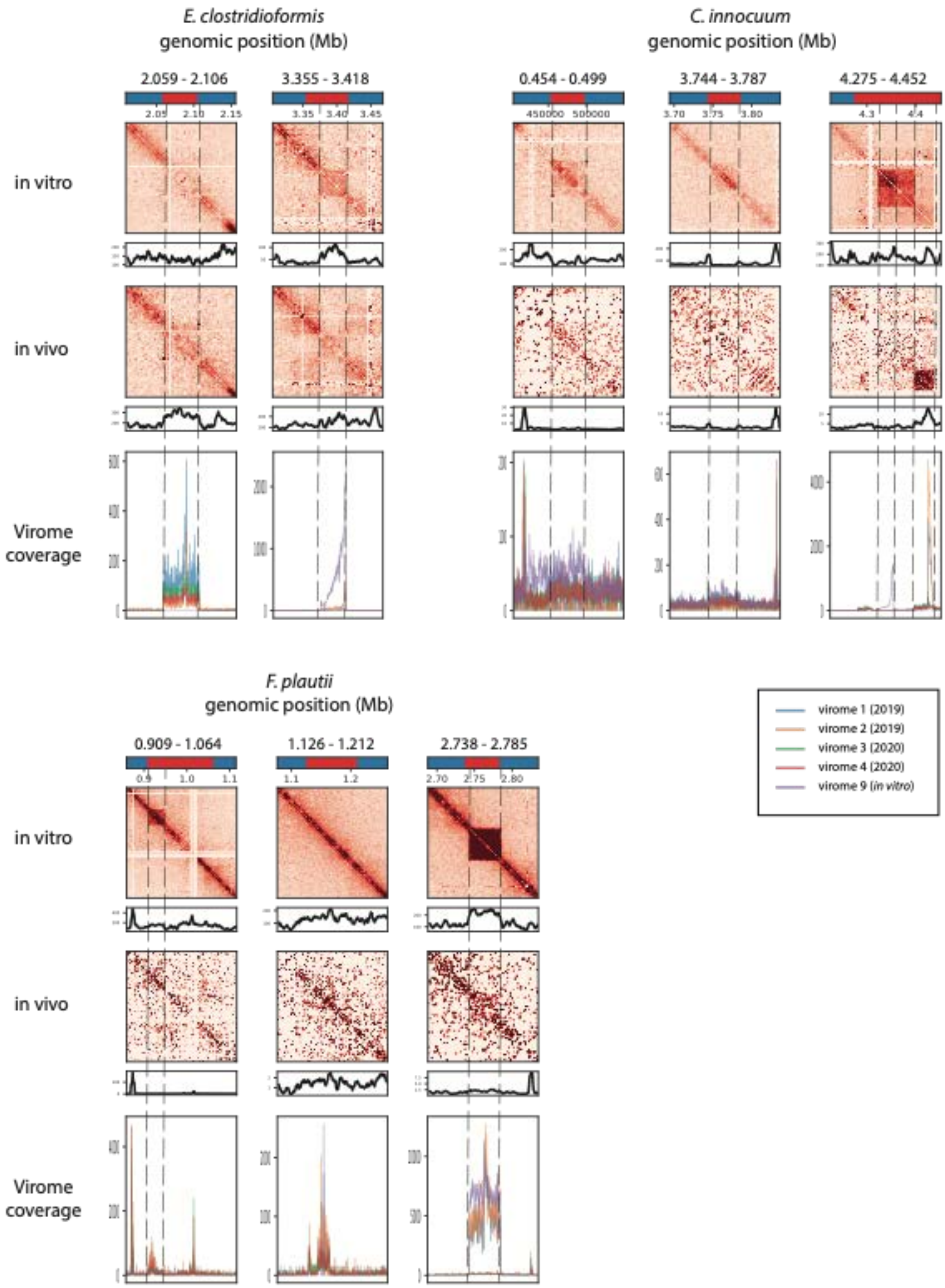

E. clostridioformis
genomic position (Mb)
2.059 - 2.106
3.355 - 3.418
C. innocuum
genomic position (Mb)
0.454 - 0.499
3.744 - 3.787
4.275 - 4.452
in vitro
in vivo
Virome
coverage
F. plautii
genomic position (Mb)
0.909 - 1.064
1.126 - 1.212
2.738 - 2.785
in vitro
in vivo
Virome
coverage
virome 1 (2019)
virome 2 (2019)
virome 3 (2020)
virome 4 (2020)
virome 9 (in vitro)

**Supplementary Figure 7: Krona representation of the Kaiju annotation of the reads not mapping on the OMM$^{12}$ strains' genomes.**

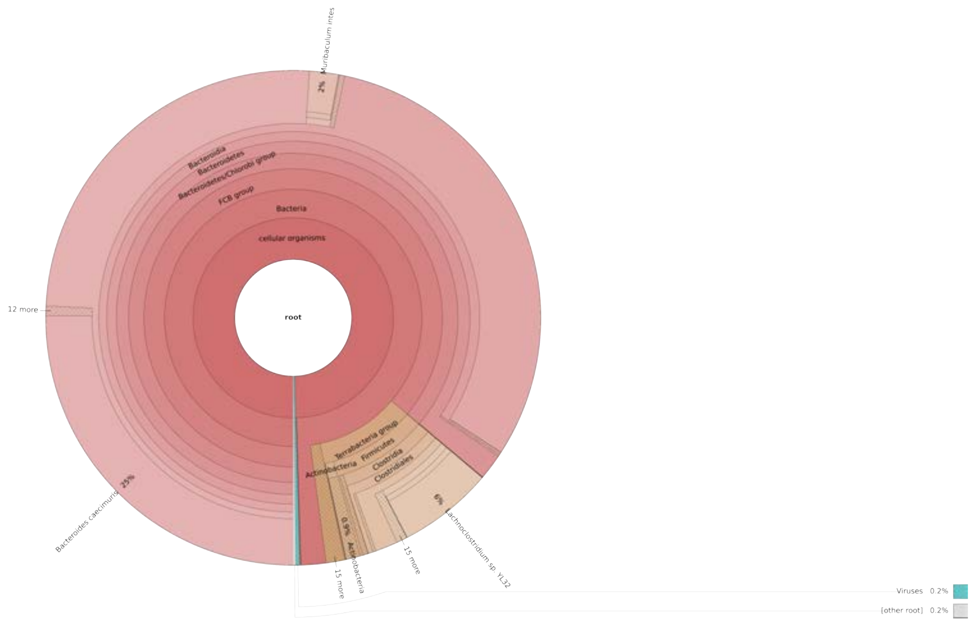

**Supplementary Figure 8: Viral clustering of the 13 induced prophages using vConctact2.** **a.** Viral cluster analysis with vConTACT2 using a gene-sharing network. The analysis was performed using genomes from the ViralRefSeq V.201 (red nodes) and CHVD (grey nodes) reference databases. Nodes for OMM[12] prophages were colored according to their respective host. **b.** Close-up view of the 13 OMM[12] inducible prophages and their direct (first level) neighbors in the network.

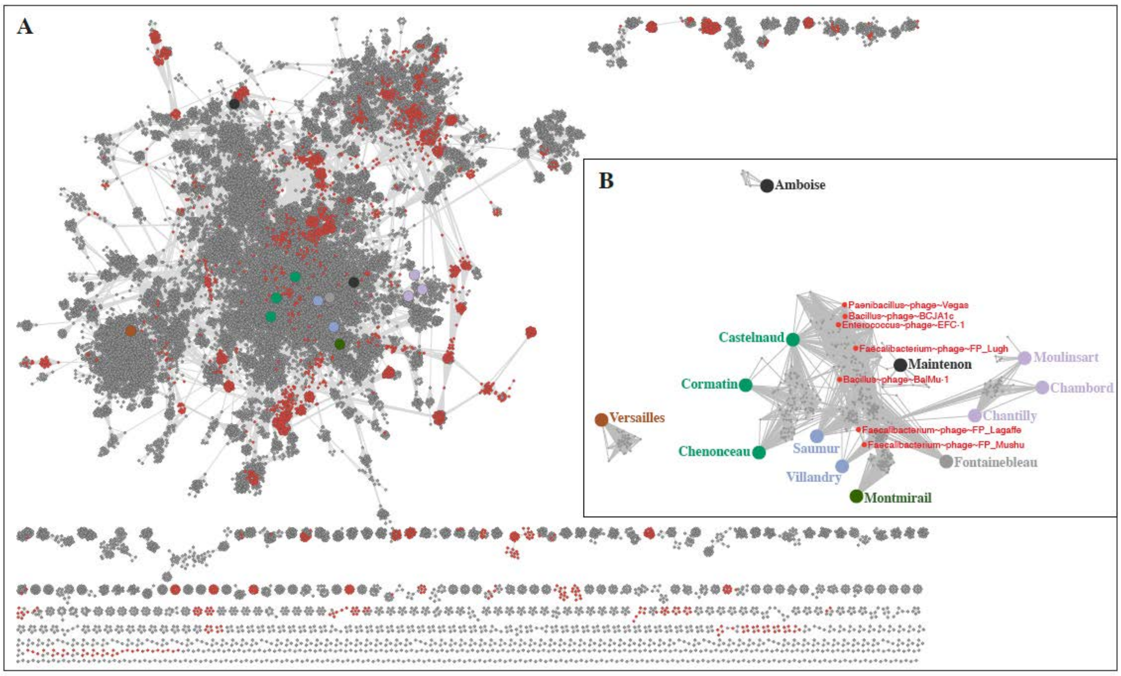

**Supplementary Table 1: Genomic libraries generated.**

a: libraries Virome_5 and Virome_6 were prepared with Accel-NGS™ 1S Plus DNA Library Kit, and thus contain accurate data for both ssDNA and dsDNA viruses. b: spike 1 was composed of $10^8$ PFU/mL of phages CLB_P1, CLB_P2, CLB_P3 and M13. c: spike 2 was composed of $10^7$ PFU/mL of phages CLB_P1, CLB_P2, CLB_P3 and M13.

| **type** | lib id | genome | species | conditions | cage | sex | sampling date | raw reads (paired-end) |
|---|---|---|---|---|---|---|---|---|
| **Hi-C *in vitro*** | OMM5 | I46 | *C.innocuum* | Lab | NA | NA | march-21 | 41 561 382 |
| | OMM5_rep | I46 | *C.innocuum* | Lab | NA | NA | July-21 | 10 688 910 |
| | OMM6 | YL58 | *B.coccoides* | Lab | NA | NA | march-21 | 21 485 845 |
| | OMM6_rep | YL58 | *B.coccoides* | Lab | NA | NA | July21 | 2 257 624 |
| | OMM7 | I48 | *B.caecimuris* | Lab | NA | NA | march-21 | 93 139 653 |
| | OMM7_rep | I48 | *B.caecimuris* | Lab | NA | NA | July-21 | 61 639 162 |
| | OMM8 | YL27 | *M.intestinale* | Lab | NA | NA | march-21 | 31 275 362 |
| | OMM8_rep | YL27 | *M.intestinale* | Lab | NA | NA | July-21 | 9 298 553 |
| | OMM9 | YL44 | *A.muciniphila* | Lab | NA | NA | march-21 | 40 820 347 |
| | OMM9_rep | YL44 | *A.muciniphila* | Lab | NA | NA | July-21 | 7 062 251 |
| | OMM10 | YL32 | *C.clostridioforme* | Lab | NA | NA | march-21 | 62 518 175 |
| | OMM10_rep | YL32 | *C.clostridioforme* | Lab | NA | NA | July-21 | 6 408 840 |
| | OMM11 | KB1 | *E.faecalis* | Lab | NA | NA | march-21 | 12 250 166 |
| | OMM11_rep | KB1 | *E.faecalis* | Lab | NA | NA | July-21 | 11 076 026 |
| | OMM12 | YL31 | *F.plautii* | Lab | NA | NA | march-21 | 10 544 208 |
| | OMM12_rep | YL31 | *F.plautii* | Lab | NA | NA | July-21 | 17 539 144 |
| | OMM13 | I49 | *L.reuteri* | Lab | NA | NA | march-21 | 9 363 639 |
| | OMM13_rep | I49 | *L.reuteri* | Lab | NA | NA | July-21 | 4 516 936 |
| | OMM14 | YL2 | *B.animalis* | Lab | NA | NA | march-21 | 7 526 155 |
| | OMM14_rep | YL2 | *B.animalis* | Lab | NA | NA | July-21 | 4 669 716 |
| | OMM15 | YL45 | *T.muris* | Lab | NA | NA | march-21 | 8 827 292 |
| | OMM15_rep | YL45 | *T.muris* | Lab | NA | NA | July-21 | 18 422 688 |
| | OMM16 | KB18 | *A.muris* | Lab | NA | NA | march-21 | 7 757 787 |
| | OMM16_rep | KB18 | *A.muris* | Lab | NA | NA | July-21 | 9 033 311 |
| **Hi-C *in vivo*** | OMM1 | mix | microbiota | microbiota | 3 | male | sept-19 | 101 182 905 |
| | OMM2 | mix | microbiota | microbiota | 4 | female | sept-19 | 115 105 053 |
| | OMM3 | mix | microbiota | microbiota | 1 | male | may-20 | 72 638 739 |
| | OMM4 | mix | microbiota | microbiota | 2 | female | may-20 | 94 694 874 |
| | Virome1 | mix | dsDNA | microbiota | 4 | female | sept-19 | 6 443 793 (2x35) |

| | | | | | | | | |
|---|---|---|---|---|---|---|---|---|
| **Virome** | Virome2 | mix | dsDNA | microbiota | 3 | male | sept-19 | 4 957 507 (2x35) |
| | Virome3 | mix | dsDNA | microbiota | 1 | male | may-20 | 108 807 445 (2x35 & 2x150) |
| | Virome4 | mix | dsDNA | microbiota | 2 | male | may-20 | 119 010 486 (2x35 & 2x150) |
| | Virome5 | mix | ssDNA[a] + spike 1[b] | microbiota | breeding | mix | march-21 | 1 056 335 (2x150) |
| | Virome6 | mix | ssDNA[a] + spike 1[b] | microbiota | breeding | mix | march-21 | 2 763 300 (2x150) |
| | Virome7 | mix | dsDNA + spike 2[c] | microbiota | breeding | mix | march-21 | 5 174 852 (2x150) |
| | Virome8 | mix | dsDNA + spike 2[c] | microbiota | breeding | mix | march-21 | 4 339 389 (2x150) |
| | Virome9 | mix (in vitro) | dsDNA | microbiota | breeding | NA | oct-21 | 6 979 488 (2x150) |

**Supplementary Table 2: Metrics of the assemblies obtained with virome reads that did not map on the OMM[12] strains.**
Both SPAdes and Megahit were used.

| | | | SPAdes | | Megahit | |
|---|---|---|---|---|---|---|
| Sample name | Sample info | Number of non mapping reads | Number of contigs | Number of contigs >5kb | Number of contigs | Number of contigs >5kb |
| #1 | 2019, 2x35 | 358 904 | 68 | 2 | 59 | 3 |
| #2 | 2019, 2x35 | 467 489 | 46 | 2 | 50 | 5 |
| #3-35 | 2020, 2x35 | 6 530 360 | 10766 | 1 | 8214 | 0 |
| #3-150 | 2020, 2x150 | 791 870 | 1611 | 2 | 2075 | 0 |
| #4-35 | 2020, 2x35 | 10 045 524 | 10241 | 0 | 5903 | 0 |
| #4-150 | 2020, 2x150 | 471 135 | 3119 | 0 | 4320 | 0 |
| merge | all above samples | 18 665 282 | 14012 | 0 | 8721 | 0 |
| merge #3 | 2020, 2x35+2x150 | 7 322 230 | 15107 | 0 | 5887 | 0 |
| merge #4 | 2020, 2x35+2x150 | 10 516 659 | 11067 | 0 | 5903 | 0 |

**Supplementary Table 3: Blast results of the contigs obtained by assembling non-mapping reads.**

| software | Contig name | Associated sample | Size | Best blast hit | Blast %coverage | Blast %id | Blast e-value |
|---|---|---|---|---|---|---|---|
| SPADES | CS1 | #1 (2019) | 24 646 | B_caecimuris | 100 | 99.90 | 0 |
| | CS2 | #1 (2019) | 22 127 | B_caecimuris | 100 | 99.78 | 0 |
| | CS3 | #2 (2019) | 53 027 | B_caecimuris | 100 | 99.8 | 0 |
| | CS4 | #2 (2019) | 7 265 | C. innocuum | 100 | 100 | 0 |
| | CS6 | #3-150 (2020, 150) | 6 639 | B_caecimuris | 100 | 99.95 | 0 |
| | CS7 | #3-150 (2020, 150) | 6 369 | B_caecimuris | 100 | 99.76 | 0 |
| | CS8 | #4-150 (2020, 150) | 5 854 | B_caecimuris | 100 | 99.79 | 0 |
| MegaHit | CM1 | #1 (2019) | 33 237 | B. caecimuris | 100 | 99.84 | 0 |
| | CM2 | #1 (2019) | 9 945 | B. caecimuris | 100 | 99.82 | 0 |
| | CM3 | #1 (2019) | 6 574 | B. caecimuris | 99 | 99.71 | 0 |
| | CM4 | #2 (2019) | 20 229 | B. caecimuris | 100 | 99.83 | 0 |
| | CM5 | #2 (2019) | 9 690 | B. caecimuris | 100 | 99.96 | 0 |
| | CM6 | #2 (2019) | 8 089 | B. caecimuris | 100 | 99.84 | 0 |
| | CM7 | #2 (2019) | 6 839 | B. caecimuris | 99 | 99.9 | 0 |
| | CM8 | #2 (2019) | 5 999 | C. innocuum | 100 | 100 | 0 |
